# Genetic load may increase or decrease with selfing depending upon the recombination environment

**DOI:** 10.1101/2021.05.20.445016

**Authors:** Shelley A Sianta, Stephan Peischl, David A Moeller, Yaniv Brandvain

**Affiliations:** University of Minnesota; University of Bern; University of Minnesota - Twin Cities

**Keywords:** Efficacy of selection, Genetic load, Mating systems, Pseudo-overdominance, Selective interference, Selfing

## Abstract

The ability of natural selection to remove deleterious mutations from a population is a function of the effective population size. Increases in selfing rate, and concomitant increases in population-level homozygosity, can increase or decrease the efficacy of selection, depending on the dominance and selection coefficients of the deleterious mutations. Most theory has focused on how (partial) selfing affects the efficacy of selection for mutations of a given dominance and fitness effect in isolation. It remains unclear how selfing affects the purging of deleterious mutations in a genome-wide context where mutations with different selection and dominance coefficients co-segregate. Here, we use computer simulations to investigate how mutation, selection and recombination interact with selfing rate to shape genome-wide patterns of genetic load. We recover various mechanisms previously described for how (partial) selfing affects the efficacy of selection against mutations of a given dominance class. However, we find that the interaction of purifying selection against mutations of different dominance classes changes with selfing rate. In particular, as outcrossing populations transition from purifying selection to pseudo-overdominance they experience a dramatic increase in the genetic load caused by additive, mildly deleterious mutations. We describe the threshold selfing rate that prevents pseudo-overdominance and decreases genetic load.

## INTRODUCTION

Populations experience an influx of deleterious mutations. How effectively natural selection removes deleterious mutations from a population depends on how clearly a mutation’s statistical effect on fitness can be seen over the genetic backgrounds in which it resides. Selection most effectively removes deleterious mutations from a large sample of independent genetic backgrounds (i.e. a large effective population size, *N*_*e*_). Deleterious mutations that are not removed make up the genetic load (i.e. a decrease in average population fitness), which can affect population viability (Lynch *et al*. 1995), patterns of introgression (Sankararaman *et al*. 2014; Kim *et al*. 2018), and further reduce selection’s efficacy in regions of low recombination (Charlesworth *et al*. 1993a; Charlesworth 1994).

Variation in the mating system, ranging quantitatively from complete outcrossing to complete self-fertilization – serves as model to better understand what modulates the efficacy of selection. Despite decades of theoretical investigation into how a population’s selfing rate affects population fitness and inbreeding depression (e.g., Charlesworth 1992; Glémin 2007), the effects of mating system on the efficacy of selection are not fully resolved in theory or empirically. We simulate whole genomes experiencing deleterious mutations with differing dominance coefficients to uncover how properties of mutations (their dominance and selection coefficients) and the genomic environment (recombination and mutation rates) interact with the mating system to affect the efficacy of selection.

The increase in individual homozygosity upon selfing modulates how mating system affects the efficacy of selection. This elevated homozygosity can decrease the efficacy of selection by decreasing the effective number of chromosomes (Pollak 1987), or increase the efficacy of selection by increasing the variance in fitness (Uyenoyama and Waller 1991). For additive mutations (*h* = 0.5), these effects cancel; the probability of fixation of such alleles is unaffected by the selfing rate (Caballero and Hill 1992; e.g., Charlesworth 1992; Glémin 2007). However, when mutation effects are less than additive effects (*h* < 0.5), the elevated homozygosity of selfers allows them to more efficiently fix advantageous (e.g., Abu Awad and Roze 2018), and remove deleterious (Charlesworth 1992) mutations. As such, partially selfing populations can more effectively “purge” recessive mutations (which are more likely to be highly deleterious than additive mutations (Crow 1993; Agrawal and Whitlock 2011; Huber *et al*. 2018) than can highly outcrossing populations (Lande and Schemske 1985). So, all else equal, the selfing rate does not change the fixation probability of a mutation with an additive effect on fitness, but the elevated homozygosity associated with selfing facilitates the purging of (partially) recessive mutations.

However, all else is not equal. Ecological, demographic, and genomic features of selfing species modulate *N*_*e*_, and thus, the efficacy of selection. On the whole, these factors tend to make selection, and particularly selection on alleles with nearly additive effects on fitness, less effective as the selfing rate (Glémin 2007; Wright *et al*. 2008; Glémin and Galtier 2012). Importantly, because recombination between homozygous sites does not generate new haplotypes, selfing decreases the effective recombination rate (Nordborg 2000). This decrease can induce Hill-Robertson interference between beneficial mutations (McVean and Charlesworth 2000) and increases the reach of classic background selection (Roze 2016). By reducing *N*_*e*_ (Charlesworth *et al*. 1993a), background selection both decreases diversity at linked neutral sites and limits selection’s ability to affect linked mutations (Charlesworth and Wright 2001). Moreover, near obligately selfing populations steadily increase their deleterious mutations by continual loss of the least loaded mutational class (a.k.a Muller’s ratchet, Charlesworth *et al*. 1993b). Thus, by increasing homozygosity, selfing allows for effective selection on (partially) recessive mutations, but by reducing the effective recombination rate selfing extends the effects of linked selection on additive mutations, decreasing the efficacy of selection.

Most theoretical studies of how mating system and linked selection interact to determine the efficacy of selection consider mutations with a single dominance level which is quite far from full recessivity (but see Arunkumar et al. (2015) for multiple mutation types with partially recessive to partially dominant mutations and Kim et al. (2018) for a dominance coefficients that continuously vary with selection coefficients). As such, our understanding of this topic is shaped by the impact of selection against rare mutations in the heterozygous state, and not dynamics that largely play out in only the homozygous state. However, purifying selection occurs in real genomes with mutations spanning a range of dominance and selection coefficients. Therefore, the impact of purging recessive mutations in highly selfing populations could mediate the consequences of the mating system on selection on linked deleterious mutations. By exposing rare deleterious recessive alleles, (partial) selfing will increase the equilibrium frequency of “unloaded” haplotypes with no such mutations (*f*_*0*_, Charlesworth et al. 1993) decreasing *N*_*e*_ and the strength of linked selection induced by recessive mutations. Thus compared to outcrossing, selfing could either reduce the impact of selective interference by effectively purging rare recessive mutations or amplify the impact of selective interference on additive mutations by decreasing the effective recombination rate. How these features interact in genomes experiencing mutations with variable selection and dominance coefficients has not been explored.

When recessive mutations arise faster than selection and recombination can remove them, obligately outcrossing populations will not be able to sustain a haplotype free of deleterious recessive mutations and will transition to a state known as pseudo-overdominance (Ohta and Kimura 1970; Gilbert *et al*. 2020). With pseudo-overdominance partially recessive deleterious alleles at different loci are maintained on complimentary haplotypes maintained by balancing selection (Charlesworth and Charlesworth 1997; Pálsson and Pamilo 1999; Charlesworth and Willis 2009). Homozygotes for any such haplotype will expose its recessive mutations, which are hidden in heterozygotes. By purging rare recessive mutations, partially selfing populations may prevent pseudo-overdominance. It also seems plausible that pseudo-overdominance could affect how effectively selection removes linked deleterious mutations. Therefore, by purging partially recessive mutations and suppressing the emergence of pseudo-overdominance, a partially selfing population may accumulate fewer linked mildly deleterious additive mutations than a highly selfing populations.

We developed a series of individual-based forward simulations to explore how selfing affects the efficacy of selection in populations experiencing both recessive and additive mutations across a wide range of parameter space. We examined how the selfing rate interacts with the action of linked selection and purging to shape the architecture of genetic load. To study how the transition from purifying selection to pseudo-overdominance interacts with the selfing rate to shape the linked load, we extend analytical models from Gilbert et al. (2020) to formally derive when background selection transitions to pseudo-overdominance as a function of the mutation, recombination and selfing rates. Our simulations recover the known effects of partial selfing on the efficacy of direct and linked selection including evidence of Mueller’s ratchet with obligate selfing (Charlesworth *et al*. 1993b) and a critical purging threshold attributable to genome-wide correlations in homozygosity under partial selfing (Lande *et al*. 1994).

We find that partial selfing can increase or decrease the efficacy of selection on linked additive mutations. When recombination rates are not very low and there is a modest input of recessive mutations selection against additive deleterious mutations is more effective in less effective in selfers than in outcrossers. On the other hand, when recombination rates are very low and recessive mutations arise frequently, selection against additive deleterious mutations is more effective in (partial) selfers than in outcrossers. Finally, we find that pseudo-overdominance further decreases the efficacy of selection and that by exposing rare recessive mutations, partial selfing prevents the transition to pseudo-overdominance.

## METHODS

We developed simulations in SLiM v3.3.2 (Haller and Messer 2019), and analytical theory to evaluate how linked selection, purging and inbreeding interact to affect the efficacy of purifying selection.

### Simulations in SLiM

#### Fixed parameters

##### Demography

All simulations consisted of 10,000 diploids over a span of 6*N* non-overlapping generations. Fitness at each locus is 1, 1 – *hs*, and 1 – *s* for genotypes homozygous for the wild-type allele, heterozygous for a deleterious mutation and homozygous for a deleterious mutation, respectively. Fitness was multiplicative across loci (i.e. the fitness of the *i*^th^ individual, *w*_*i*_ = ∏*w*_*ij*_ = ∏(1 – *h*_*ij*_ *s*_*ij*_), where *j* indexes the locus).

##### Genome size and structure

Genomes consisted of six 7.5Mb chromosomes, as in Gilbert et al. (2020), with uniforn recombination rates across each chromosome, and free recombination among chromosomes. We modeled a uniform genome structure, in which mutation each mutation type (see below) was independent of genomic position.

##### Mutational effects

SLiM simulates genomes composed of specific “mutation types,” each characterized by a fixed dominance coefficient, *h*, and a (distribution of) selection coefficient(s), *s*. All simulations included four deleterious mutation types – one fully to partially recessive mutation (0 ≤ *h* < 0.5) type, three additive mutation (*h* = 0.5) types, and no neutral or beneficial mutation types. As such the genome-wide mutation rate, *U*, represented the deleterious genome-wide mutation rate, *U*_*del*_. Each of the four mutation types contributed equally to *U*_*del*_ such that the total additive deleterious mutation rate was three times that of the recessive deleterious mutation rate.

##### Selection coefficients for mildly deleterious, additive mutations

We chose selection coefficients of the three additive mutation types to slightly exceed the nearly neutral boundary (4*N*_*e*_*s* > 1), which differentiates where natural selection can and cannot effectively remove deleterious mutations (Kondrashov 1995) because this should expose differences in the efficacy of selection against additive load in selfers and outcrossers. The selection coefficients for these three mutation types were *s* = 0.0005 (4*Ns* = 20), *s* = 0.00025 (4*Ns*=10) to *s* = 0.00005 (4*Ns*=2). Given that *N*_*e*_ is likely less than the census population size (*N*) in all multilocus simulations, we assume the 4*N*_*e*_*s* values will be less than the 4*Ns* values listed above. We chose these fixed selection coefficients, rather than the more biologically realistic distribution of fitness effects, because they provide theoretical insight into when and how selection becomes less effective.

#### Variable model parameters

We investigated all factorial combinations of five variables: (1) selfing rate, (2) deleterious mutation rate, (3) recombination rate, (4) fitness cost of strongly deleterious recessive mutations (*s*_*recessive*_) and (5) recessivity of strongly deleterious mutations (*h*_*recessive*_), with ten replicates for each parameter combination.

##### Selfing rate

Selfing rates ranged from obligate outcrossing (α = 0) to near obligate selfing (α = 0.99) and values between (α = 0.05, 0.1, 0.25, 0.5, 0.75, or 0.9), allowing us to evaluate both mixed maters as well as predominant selfers or outcrossers. Because we do not model an evolutionary transition between selfing rates, we do not consider the initial purging, or lack thereof, of the load required for the transition to selfing (see Wang *et al*. 1999; Bataillon and Kirkpatrick 2000; Waller 2021).

##### Deleterious mutation rate

We varied the genome-wide deleterious mutation rate (*U*_*del*_ = *μ*_*del*_ x genome size), as *U*_*del*_ modulates background selection (Charlesworth *et al*. 1993a; Kamran-Disfani and Agrawal 2014). We chose *U*_*del*_ values of 0.04, 0.16, and 0.48 to span a range of *U*_*del*_ values estimated from multicellular eukaryotes (Schultz *et al*. 1999; Willis 1999; Cutter and Payseur 2003; Haag-Liautard *et al*. 2007; Lynch 2010; Slotte 2014).

##### Recombination rate

To probe how linked selection interacts with the selfing rate to modulate the efficacy of selection, we varied the recombination rate. We report this as the Relative Recombination Rate (RRR) – the per-base-pair recombination rate divided by the per-base-pair mutation rate. We examined RRR values of 0.01, 0.1, 1 and 10, corresponding to per-base-pair recombination rates ranging from 8.89 × 10^−12^ to 1.07 × 10^−7^ across all mutation rates (see Table S1 for per-base pair mutation and recombination rates for each parameter combination).

##### Selection and recessivity of (partially) recessive deleterious mutations

We varied the intensity of selection against strongly deleterious (partially) recessive mutations from *s*_*recessive*_ = 0.015, *s*_*recessive*_ = 0.3, and *s*_*recessive*_ = 0.9. These values all prevent the chance fixation of such mutations, but the greater the selection coefficient the more rapidly a mutation is removed. These mutations could be nearly additive (*h*_*recessive*_=0.25), partially recessive (*h*_*recessive*_ = 0.1), or fully recessive (*h*_*recessive*_ = 0).

#### Quantifying the consequences of selection

##### Quantifying the efficacy of selection

We initially examined the load of recessive and additive mutations separately. We assessed the efficacy of selection both in terms of prevalence (the average number of mutations per individual) and the overall fitness consequences for each of the four mutation types. Throughout our results, all discussions of dominance and/or additively refer to the mode of gene action (a parameter we specify in our model), not the components of genetic variance.

Prevalence is meant to capture common genomic summaries of the number of derived deleterious mutations, while the genetic load is meant to quantify the fitness consequences of these mutations (Lohmueller 2014; Do *et al*. 2015). Because the degree of homozygosity increases as a function of selfing rate, we expected that the translation of prevalence to fitness of recessive mutations would vary across selfing rates. By contrast, additive mutations are expressed to some extent across all populations and thus patterns of prevalence across selfing rates should translate more directly to patterns of overall fitness. We also report the mean population fitness, including both additive and recessive mutations.

For computational efficiency, SLiM only tracks segregating mutations; consequently, population fitness outputted by SLiM excludes the effects of fixed mutations. We therefore developed custom R scripts to calculate both mutation prevalence and population mean fitness from both segregating and fixed mutations in all simulations.

##### Summarizing neutral genetic variation

Pseudo-overdominance can leave a genomic signature of elevated diversity at linked neutral sites (Gilbert *et al*. 2020). This signature arises when heterozygotes at neutral sites appear to have higher fitness than homozygotes – a phenomenon known as associative overdominance (Frydenberg 1963). We quantified two genomic signatures of associative overdominance as a proxy for whether pseudo-overdominance occurred, namely increased neutral diversity (π) and an intermediate-frequency-skewed unfolded allele frequency spectrum (AFS), in eight replicate runs per simulation. Because explicitly modeling neutral mutations is computationally burdensome, we used the tree sequence recording function within SLiM (Haller *et al*. 2019). We subsequently overlaid neutral mutations on each tree sequence at a mutation rate of μ = 1e-7 in msprime (Kelleher *et al*. 2016), sampled one genome from 200 individuals, and calculated π and the AFS with the 200 samples using tskit (https://tskit.dev/tskit/docs/stable/).

Numerous processes – including true overdominance (Ohta and Kimura 1971), selection against recurrent recessive mutations (Zhao and Charlesworth 2016; Becher *et al*. 2020), genome-wide selection against homozygosity when genotypes are correlated at unlinked loci (a.k.a. identity disequilibrium, Charlesworth 1991), and pseudo-overdominance (Gilbert *et al*. 2020) can generate a pattern of associative overdominance. However, because our simulations did not allow for recurrent mutation nor classical overdominance, and because our selection coefficients can lead to pseudo-overdominance but not to associative overdominance (i.e., when *Ns* > 1, Becher et al. (2020)), these explanations are not relevant for this study. Rather only pseudo-overdominance and genome-wide selection against homozygosity could potentially explain signatures of associative overdominance in our simulations.

### Analytical Model for the transition to pseudo-overdominance

To analytically derive how partial selfing prevents the transition from background selection to pseudo-overdominance, we extended the multi-locus model of Gilbert et al. (2020) to include selfing (Appendix). We considered *n* biallelic loci and denoted wild-type and derived alleles at locus i by a_*i*_ and A_*i*_, respectively. Derived alleles were deleterious and fully recessive with fitness 1− s when homozygous and 1 otherwise, with multiplicative fitness effects across loci.

We derived the frequency of the zero-mutation haplotype at mutation-selection balance. When this frequency approaches zero, we expect a transition from background selection to pseudo-overdominance. We assumed that haplotypes with more than one deleterious mutation are vanishingly rare at equilibrium and hence ignored them, meaning that any genotype can be polymorphic at most at two deleterious loci. This assumption is quite different than what we find in our SLiMulations, but provides a reasonable guide to our major qualitative results.

We used *p*_*i*_, *i* = 1,…, *n*, to denote the frequency of the haplotype carrying a derived deleterious allele at locus *i*, and *p*_O_ the frequency of the haplotype without any deleterious mutations. Loci were equidistantly distributed over a region of length *r* cM, such that the recombination rate between adjacent loci is *r*/(*n* − 1). We followed the frequencies of four genotypic classes: genotypes without any deleterious mutations (*p*_00_), genotypes heterozygous at exactly one locus 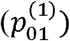, genotypes heterozygous at exactly two loci 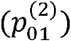 and genotypes homozygous for the derived allele at exactly one locus (*p*_11_). Fitnesses of these genotype-classes are 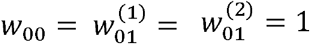 and *w* _11_ =1 − *s*, so mean fitness is 1- *p*_11_ *s*. In the Appendix we derived equations for how the frequencies of these genotypic classes change over time. We then derived the exact equilibrium frequency of the zero-mutation haplotype could be readily determined in the absence of recombination and approximated it for weak recombination and selfing rates in eqs. (1) and (2) in the Appendix.

## RESULTS

### Selfing rate can increase or decrease the genetic load in multilocus simulations

An increase in the selfing rate either slightly or severely reduces the frequency of recessive deleterious mutations, and can increase, decrease, or not change the burden of mildly deleterious additive mutations. We explore these alternative outcomes, and what modulates them, below: first for recessive strongly deleterious (*Ns* > 150) recessive mutations, then additive mildly deleterious (*Ns* < 5) mutations, and finally for overall population fitness.

### Selfing prevents the accumulation of recessive mutations and impedes the transition to pseudo-overdominance

#### Complete recessivity and relatively high recombination rates

Highly selfing populations maintain a lower prevalence and frequency of recessive mutations than do outcrossing populations (Figure 1). With high recombination rates (RRR ≥ 1), the number of recessive deleterious mutations per individual generally declined with the selfing rate. This decay is rapid and steep when mutations are strongly deleterious. That is, populations with mixed mating systems effectively purge extremely deleterious mutations (*s*_*recessive*_= 0.3 or *s*_*recessive*_= 0.9) but maintain a larger number of less deleterious mutations (*s*_*recessive*_ = 0.015, Figure 1A). These results are consistent with both our analytical derivations and those by Roze and Rousset (2004) for the frequency of a recessive deleterious mutation at a single locus as a function of selfing rate (see equation A1 and compare Figures 1B and Figure A1 in the Appendix), which is expected as loci should be more or less independent at high relative recombination rates.

**Figure 1:**
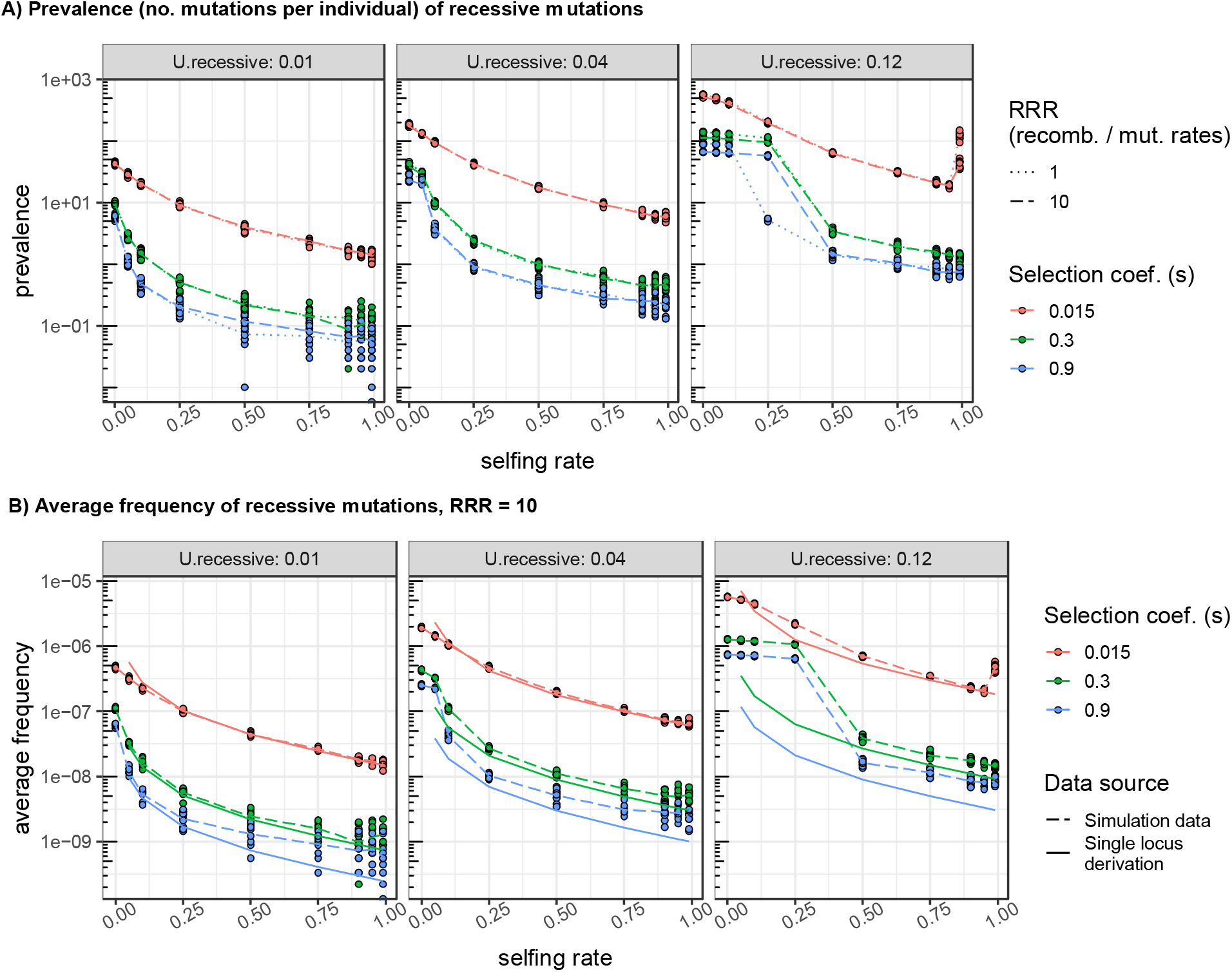
Purging dynamics of recessive mutations when per base-pair recombination rates are equal to (relative recombination rate, RRR, = 1) or greater than (RRR >1) per base-pair deleterious mutation rates. (A) Prevalence (i.e. the mean number of mutations per diploid genome) of recessive mutations. The tick marks on the y-axis highlight the log_10_ scale. (B) Results from multilocus simulations tend to fit analytical expectations derived from single locus models better with weaker selection coefficients and lower mutation rates than with large selection coefficients and higher mutation rates. This likely reflects cases in which identity disequilibria generated by partial selfing hinders the purging process (at *U*_*recessive*_ = 0.12 and *s*_*recessive*_ = 0.3 and 0.9), and follows from the assumption made in our analytical derivation that mutation rates are much smaller than selection coefficients.

The frequency of recessive mutations drops dramatically between selfing rates of 0.25 and 0.5 when the mutation rate is high and mutations are quite severe (*U*_*recessive*_ = 0.12, *s*_*recessive*_ = 0.9 or *s*_*recessive*_ = 0.3), and shows a similarly dreastic shift as the selfing rate increases from 0.05 and 0.10 for the most damaging mutations at intermediate mutation rates (*U*_*recessive*_ = 0.04 *s*_*recessive*_ = 0.9, Figure 1B). We interpret these shifts as a “purging threshold” which differentiates selfing rates that can and cannot effectively purge their recessive load (Lande *et al*. 1994). Given the high recombination rate in these simulations (this effect is strongest with an RRR of 10), we attribute this purging threshold – which exceeds single locus expectations – to the near-lethal inbreeding depression that occurs when selfed individuals in partially selfing populations expose numerous unlinked recessive mutations. This correlation in individual homozygosity across unlinked loci is a form of identity disequilibrium (Weir and Cockerham 1973), is greatest in partially selfing populations, and can hinder the purging of the recessive load because selfing events do not generate living offspring (Lande *et al*. 1994; Kelly 2007). The fit between multi-locus simulation results and analytical single locus derivations is qualitatively tighter with weaker selection, lower mutation rates and lower selfing rates, likely because multilocus interactions due to linked selection in predominant selfers are greater with stronger selection (Charlesworth *et al*. 1993a) and because analytical derivations assume weak selection and low mutation (i.e., *u* << *s* << 1).

The other deviation between single locus theory and multilocus simulation is the increased prevalence and frequency of recessive mutations with more modest effects on fitness at very high selfing rates when mutation is high (Figure 1). Given the very low effective recombination rate in highly selfing populations, this spike presumably reflects the action of Mueller’s ratchet (Charlesworth *et al*. 1993b).

#### Complete recessivity and relatively low recombination rates

We also observe a drastic drop in the frequency of recessive mutations at a critical selfing rate, which we again call a purging threshold, when mutation rates exceed recombination rates (RRR < 1, Figure 2A). This purging threshold has a much greater magnitude than that seen at high relative recombination (see right panel in Figure S1A at *s*_*recessive*_ = 0.3 and 0.9). Additionally, this dramatic shift is observed across a broader range of absolute mutation rates, and with much weaker selection coefficients than we observed with relatively high recombination rates (Figure S1A).

**Figure 2:**
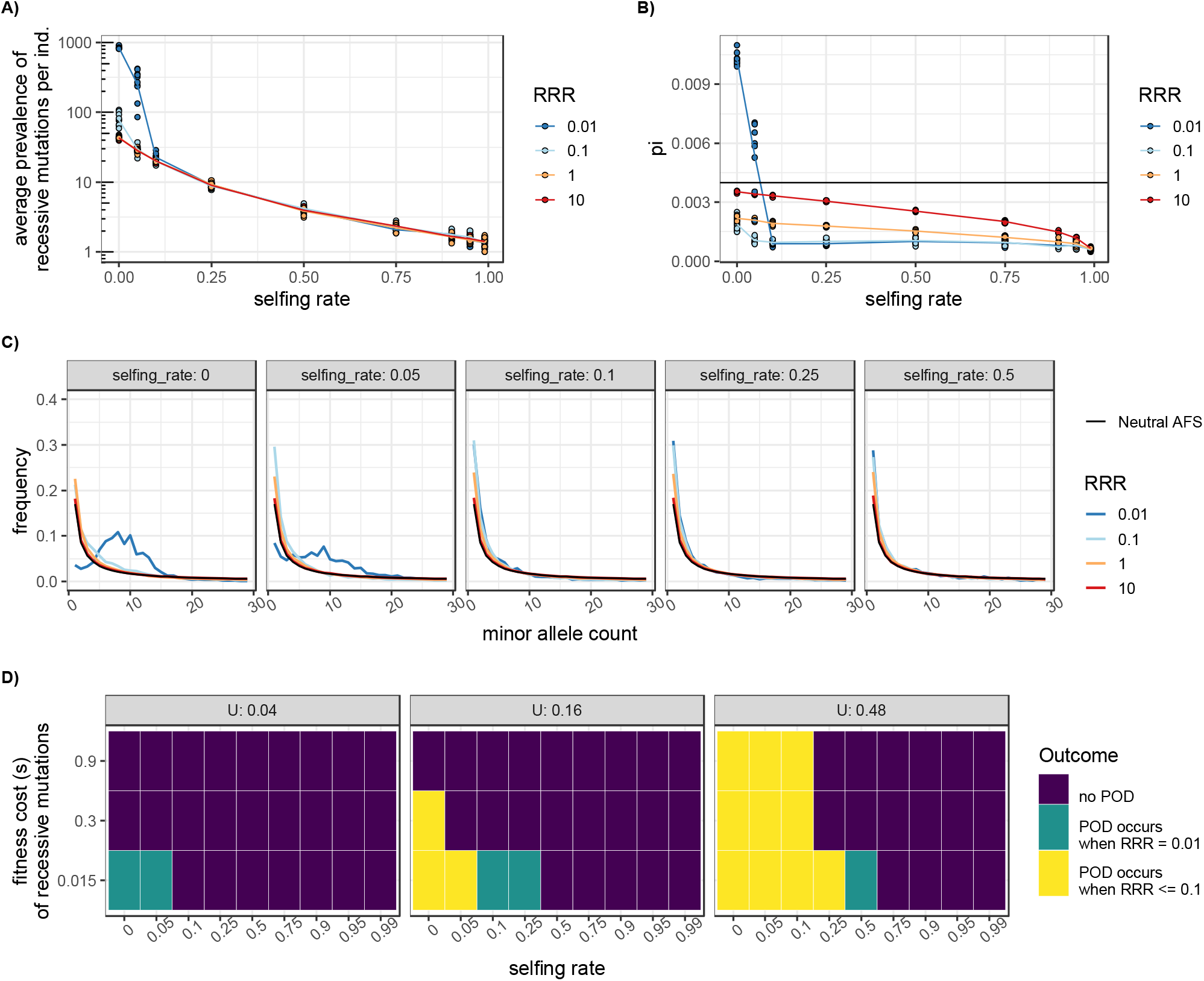
Pseudo-overdominance (POD) occurs in low recombination environments (relative recombination rate, RRR, < 1). (A) POD leads to a spike in the prevalence of recessive mutations in predominantly outcrossing populations. Points are simulation replicates and lines connect mean values. Genome-wide deleterious mutation rate *U*_*del*_ = 0.04, *s*_*recessive*_= 0.015, and *h*_*recessive*_ = 0. (B) POD also leads to a spike in neutral diversity, driven by heterozygosity at linked neutral sites. *U*_*del*_ = 0.04, *s*_*recessive*_= 0.015, and *h*_*recessive*_ = 0. Expected neutral diversity (4Nμ) is shown by the black horizontal line. (C) Allele frequency spectra (AFS) at a subset of selfing rates (different facets) for *U*_*del*_ = 0.04, *s*_*recessive*_= 0.015, and *h*_*recessive*_ = 0. POD shifts the AFS to more intermediate frequency alleles. Mean AFS are in bold lines, and individual simulation replicates are in faint lines. Black lines correspond to the neutral AFS. (D) Outcome plot of POD occurrence when *h*_*recessive*_ = 0. Green blocks indicate POD occurs at only the lowest relative recombination rate (RRR; RRR = 0.01); yellow blocks indicate POD occurs at the two lowest RRR (0.01, 0.1). Data in panels A-C correspond to the bottom row of the *U*_*del*_ = 0.04 outcome plot.

We attribute this drastic difference in the recessive load in populations above and below the purging threshold in low RRR simulations to a transition from classic purifying selection to pseudo-overdominance in primarily outcrossing populations. The structuring of pseudo-overdominant haplotypes by complementary sets of deleterious recessive mutations in repulsion is clearly visible in samples of genomes from populations below this purging threshold and is not observed in populations which can more effectively purge (Figure S2). Consistent with this explanation, simulation runs that appear to display a shift to pseudo-overdominance are also associated with elevated genetic diversity (π) at linked neutral sites (Figure 2B, Figure S3A) and a shift towards more high-frequency alleles (Figure 2C, Figure S4), two consequences of pseudo-overdominance (Gilbert *et al*. 2020). This increased neutral diversity is in contrast with classic background selection, which reduces linked neutral diversity. We see the signature of background selection by the reduction in π with a reduction in the relative recombination rate (Figure S3).

Partial selfing can prevent the shift to pseudo-overdominance by exposing the deleterious effects of rare recessive mutations. The amount of selfing required to prevent the emergence of pseudo-overdominance depends on how rapidly deleterious recessive mutations can be removed by selection before mutation-free haplotypes are eliminated. As such, less selfing is required to prevent the emergence of pseudo-overdominance when mutations have more severe fitness consequences, while more selfing is required to prevent the emergence of pseudo-overdominance when the mutation rate is high (Figure 2D). As expected, pseudo-overdominance evolves more readily as the recombination rate decreases (compare RRR of 0.01 to RRR of 0.10 in Figure 2D). Overall, we observe signatures of pseudo-overdominance when selfing is rare, recombination is infrequent, and *U*_*del*_*/s* > 0.5.

Once pseudo-overdominant haplotypes form, homozygote fitness plummets. With partial selfing (or more recombination) fitness due to recessive mutations drops further, as recombination creates more non-complementary haplotypes (the recombination load) and/or as selfing exposes more haplotypes in the homozygous state (the segregation load, Figure S5).

#### Partial recessivity across recombination rates

Because partial recessivity increases the capacity for selection to remove partially recessive mutations in predominantly outcrossing populations, we tested the possibility that intermediate dominance coefficients (*h*_*recessive*_= 0.1 and *h*_*recessive*_= 0.25) could prevent the emergence of pseudo-overdominance. Like Gilbert et al. (2020), we find that, although partial recessivity substantially decreases the parameter space under which pseudo-overdominance occurs, it does not prevent it altogether (Figure S6-S9). When *h*_*recessive*_= 0.1, pseudo-overdominance occurs when *s*_*recessive*_ is relatively modest (*s*_*recessive*_= 0.015; *Nhs* = 15) and mutation rates are high (U_del_=0.48) at selfing rates of 0, 0.05, 0.1 and 0.25. By contrast when deleterious mutations are more damaging, partial recessivity more effectively prevents the emergence of pseudo-overdominance because these mutations are effectively removed when heterozygous. When *h*_*recessive*_ = 0.25, partially recessive mutations accumulate in predominantly outcrossing populations at the highest mutation rate and lowest selection coefficient (*s*_*recessive*_= 0.015; *Nhs* = 37.5; Figure S1-C), but the accumulation does not cause the switch to pseudo-overdominance, i.e., there is no increase in diversity (Figure S3-C) nor a shift in the allele frequency spectrum (Figure S9). Thus, while partial recessivity limits the extent of pseudo-overdominance, it does not eliminate it.

In the remainder of the considered parameter space (i.e. lower mutation rates, higher relative recombination rates, and/or more damaging mutations), increasing the dominance coefficient of recessive mutations decreases prevalence (number of mutations per individual) in the primarily outcrossing populations, as expected. Prevalence still decreases with selfing rate, but the absolute difference in prevalence between outcrossers and selfers diminishes (Figure S7).

### An analytical model for the transition to pseudo-overdominance with partial selfing

Our SLiMulations revealed that (partial) selfing can prevent the emergence of pseudo-overdominance. However, without a computationally intense search of parameters space, it does not allow us to quantitatively characterize this threshold. To do so, we present results of our analytical model which approximates the frequency of the unloaded haplotype (i.e. the haplotype with no derived deleterious mutations) under the assumption of weak selfing and weak recombination (equation (2) in the Appendix). Roughly speaking, this assumes that *F, r, u* < 1/*N* < *s*, but the exact conditions for when our approximation will become unreliable are difficult to derive (see Appendix for more details). Reassuringly, when reduced to a single locus, our model recovers the results of Roze and Rousset (2004). Because comparable two-locus results have not been derived previously, we check our approximation against results obtained by numerical iteration of the difference equation (see Appendix Figure A1 and A2). Importantly, the critical selfing rate for loss of the zero-mutation haplotype in two-locus simulations is accurately predicted by our model (see Appendix Figure A2).

We find that the frequency of the unloaded haplotype approaches zero (i.e. when we expect pseudo-overdominance) when selection coefficients are small, selfing is rare, and recombination rates are low (Figure 3). When recombination is much rarer than mutations to recessive mutations (RRR = 0.01, Figure 3A), frequent selfing is required to prevent the transition to pseudo-overdominance when selection coefficients are small. By contrast, lower selfing rates can prevent the transition to pseudo-overdominance as the relative recombination rate increases (RRR = 0.1, Figure 3B). This synergistic effect of recombination and selfing on the efficacy of purging the recessive load and preventing pseudo-overdominance is mostly observed for small selection coefficients, simply because pseudo-overdominance is unlikely if selection is strong and hence there is no opportunity for recombination to prevent it. The synergistic effect of recombination and selfing is quite general in our model (Appendix Figure A3) and is consistent with results from our whole genome SLiMulations (Figures 2D and 3).

**Figure 3:**
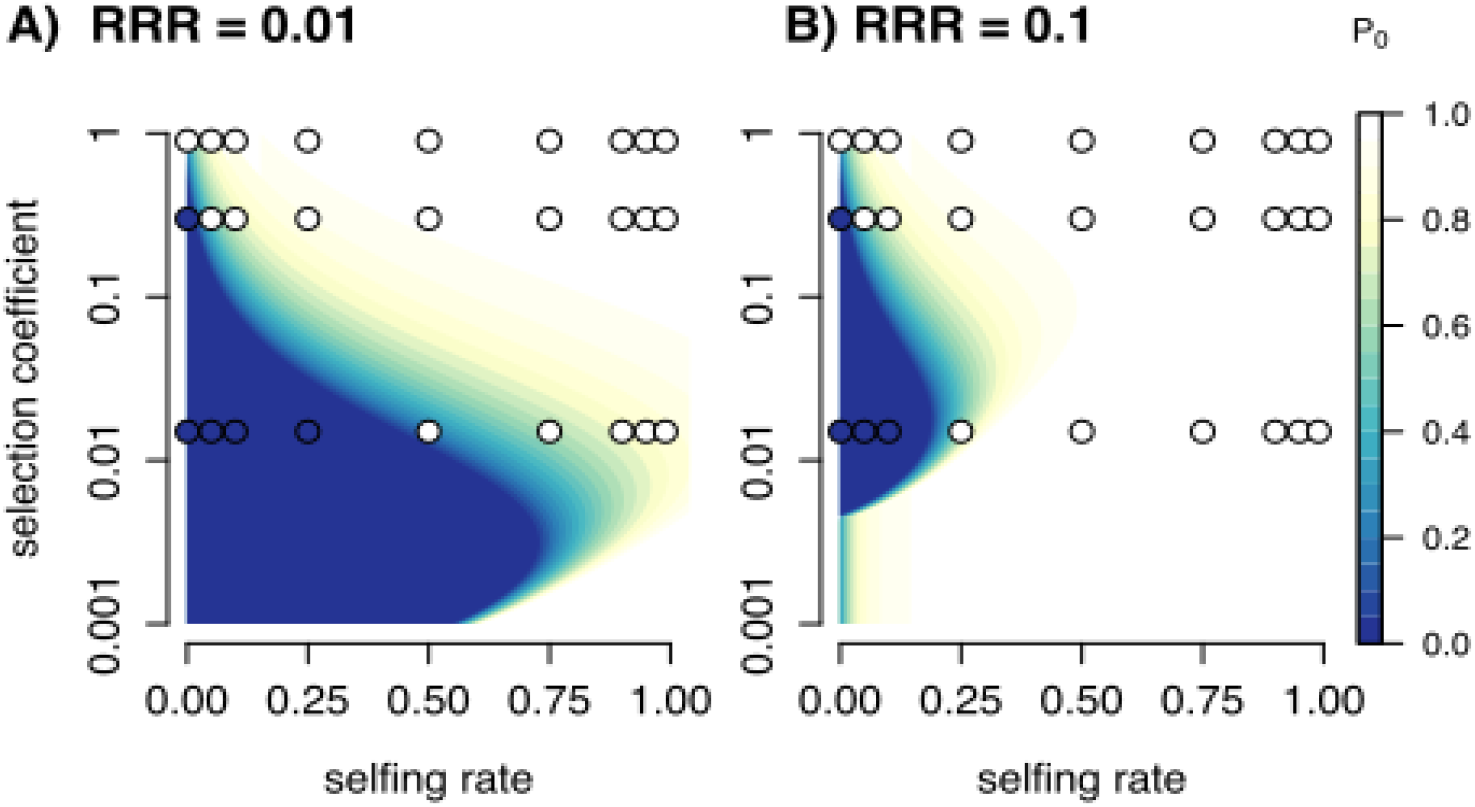
Qualitative comparison between analytical predictions and simulation results. Shaded areas indicate the analytical predictions for pseudo-overdominance (POD), specifically the frequency of the zero-mutation haplotype (P_0_), based on equation (2; see Appendix) for various recombination rates. For the analytical model, we assumed a total of loci equidistantly spaced on a chromosome with a total mutation rate of *U*_*del*_ = 0.005 and relative recombination rate RRR = 0.01 (A) and 0.1 (B). Circles show results from simulations when *U*_*del*_ = 0.16: filled circles indicate simulations where we observed POD and white circles indicate no POD.

Our analytical predictions are qualitatively consistent with the SLiMulation results (Figure 3). However, because our approximation is not tailored to direct comparison with genome-wide simulations of thousands of loci in regions of low recombination, the parameters used for the comparison between simulations and the analytical model in Figure 3 are not directly comparable (specifically, we assumed a much smaller number of loci and larger per locus mutation rate in the analytical model as compared to the simulations).

### The additive load does not necessarily increase with selfing

When modelling both additive and recessive deleterious mutations, an increase in the selfing rate can have no effect, increase, or decrease the genetic load (Figure 4). These different outcomes are determined by (1) the strength of selective interference induced by (partially) recessive mutations, and (2) whether predominantly outcrossing populations experience pseudo-overdominance.

**Figure 4:**
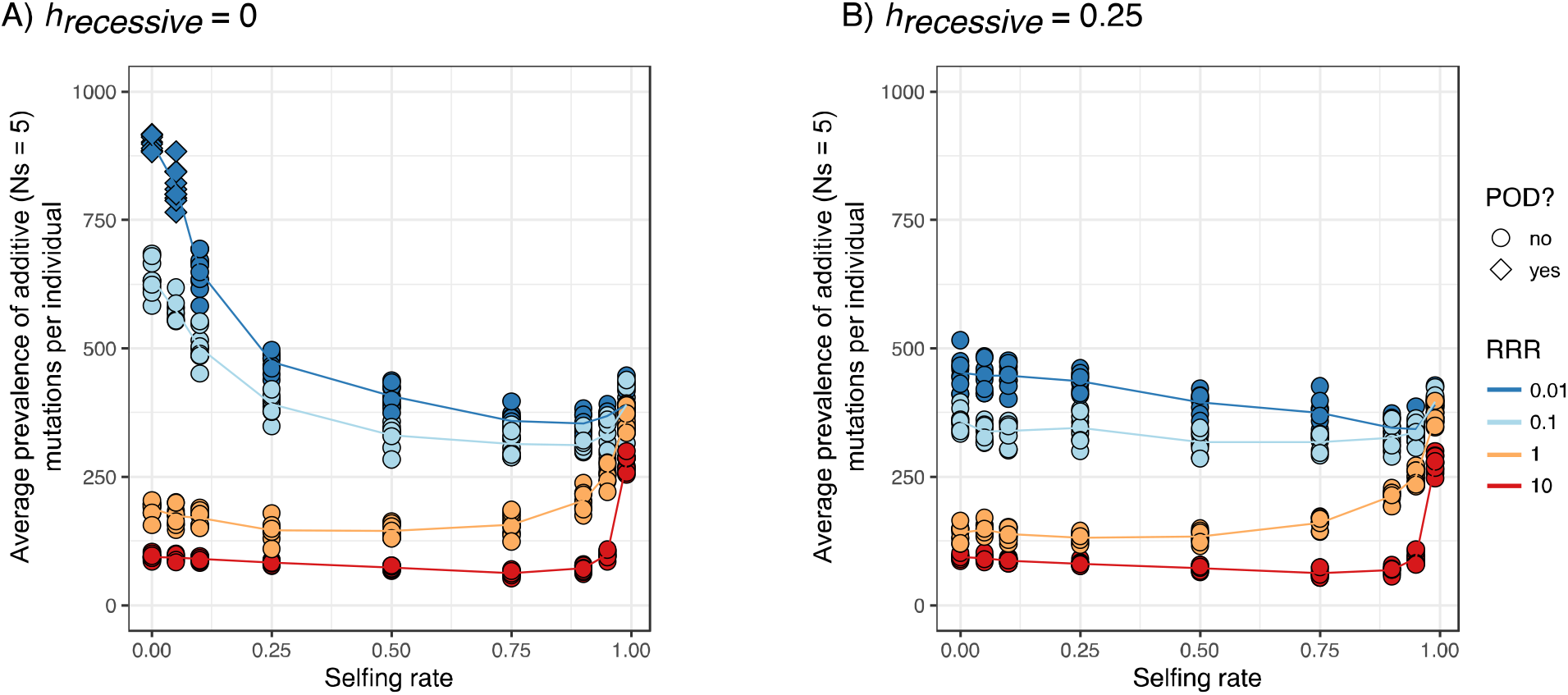
The accumulation of additive mutations as a function of selfing rate is heavily influenced by the relative recombination rate (RRR) and whether pseudo-overdominance (POD) occurs. (A) and (B) contrast simulations where the recessive load is fully recessive (A) and partially recessive (B). *U*_*del*_ = 0.04 and *s*_*recessive*_*=* 0.015.

To demonstrate the effects of selection on partially versus completely recessive mutations on selection against additive mutations, we contrast simulations with fully recessive mutations (*h*_*recessive*_= 0) to simulations with partially recessive mutations (*h*_*recessive*_= 0.25). Throughout, we present the prevalence of the most deleterious additive mutation type (*s*_*additive*_ = 0.0005, *Ns* = 5), as results are qualitatively similar across the three mutation additive mutation types (Figures S10-12).

#### Selfers evolve a larger additive load than outcrossers when recombination rates are high

Predominant selfers accumulate a higher prevalence of additive mutations in high recombination environments than do predominant outcrossers (recombination rate is greater than (red) or equal to (yellow) the mutation rate, Figures 4, S10-12). At the highest relative recombination rates (RRR = 10), near obligate selfing (selfing rates greater than 0.95) is required for an increase in the prevalence of additive mutations, as the local effective recombination rate is sufficiently large to allow most mutations to escape selective interference otherwise. At the highest recombination rate, the additive load subtly decreased with the selfing rate until the selfing rate became high enough to experience selective interference (∼ 0.75-0.9). We revisit this result, which was also observed in Roze (2015), in our low recombination rate results below. We find a similar pattern when the recombination rate equals the mutation rate, however, in this case, the additive load begins to increase at a lower selfing rate. At both of these recombination rates, the prevalence of additive mutations is very similar for cases in which strongly deleterious mutations are fully or partially recessive. With high recombination rates, the recessive load is low and pseudo-overdominance never occurs (compare orange and red lines in Figure 4A i.e. *h* = 0.00 to those in Fig 4B i.e. *h* = 0.25).

When recessive mutations have modest effects on fitness (*s*_*recessive*_ = 0.015) and the absolute mutation rate is large, there is a greater uptick of the prevalence of additive mutations in primarily selfing populations, regardless of recombination rate (Fig S10 A-C, top row). This pattern corresponds to the increase in the prevalence of recessive mutations at the same parameter values (e.g. right panel of Figure 1A), which we attribute to Mueller’s ratchet (Charlesworth *et al*. 1993b).

#### With low recombination, the additive load in selfers is equal to or smaller than that of outcrossers

When recombination is rarer than mutation, purging of recessive mutations under partial selfing can increase the efficacy of selection against linked mildly deleterious additive mutations. With a relative recombination rate of 0.1 (light blue in Figure 4A), the prevalence of additive mutations decreases as populations transition from obligate outcrossing to predominant selfing, and the pattern becomes more dramatic as the relative recombination rate decreases further (RRR = 0.01, dark blue). As outcrossing populations transition from purifying selection to pseudo-overdominance (diamonds in Figure 4), they accumulate many more deleterious additive mutations than (partially) selfing populations.

Figures 4A vs B contrasts the same set of parameter conditions (*U*_*del*_ = 0.04 and *s*_*recessive*_= 0.015) between simulations with fully recessive (*h =* 0.00, Figure 4A) and intermediate recessive (*h* = 0.25, Figure 4B) mutations. Strongly recessive mutations (but not mutations with intermediate dominance coefficients) generate a dramatic uptick in the prevalence of mildly deleterious additive mutations at very high outcrossing rates and very low recombination rates. This dramatic result does not require complete recessivity – simulations with *h* = 0.1 can also generate a similarly dramatic spike at high outcrossing rates (Figure S10). However, this uptick is not observed at comparable simulations for higher levels of dominance (*h* = 0.25, Figures 4, S10).

#### Pseudo-overdominance decreases the efficacy of selection on linked mutations with additive effects

We propose that pseudo-overdominance limits the efficacy of selection by effectively subdividing the population into haplotypic classes of complementary recessive mutations. We show that when an additive deleterious (or beneficial) mutation falls on a haplotype maintained at equilibrium by pseudo-overdominance, selection against (or for) the new mutation will be limited by the recessive load at linked sites. Specifically, the Appendix shows that in a two-locus model for pseudo-overdominance the efficacy of selection is reduced by a factor of 1-*s*/2 for outcrossing populations, where *s* is the fitness effect of recessive mutations. Intuitively speaking, in our two-locus model a new additive mutation will have a 50% chance to be in a beneficial heterozygous genotype where the recessive load is masked, effectively reducing the strength of selection against (or for) the additive mutation by (1-*s*) in half of the genotypes. These results resemble Assaf et al.’s (2015) “staggered sweep” model, in which the spread of adaptive mutations is slowed by the exposure of linked recessive mutations that occurs when they become common.

### The effect of mating system on mean population fitness

When recombination rates are high relative to mutation rates, mean population fitness is generally greatest in outcrossers and lowest in selfers (Figure 5, Figure S13), reflecting the elevated additive load accumulated by selfers (Figure 4A). Exceptions are at the highest recombination rates, when fitness is maximized in high partial selfers. By contrast when recombination rates are lower than mutation rates, mean population fitness either does not vary with selfing rate (in the absence of pseudo-overdominance), or increases with selfing rate (in the presence of pseudo-overdominance). The effect of pseudo-overdominance on overall fitness is primarily due to its effect on the prevalence of additive mutations.

**Figure 5:**
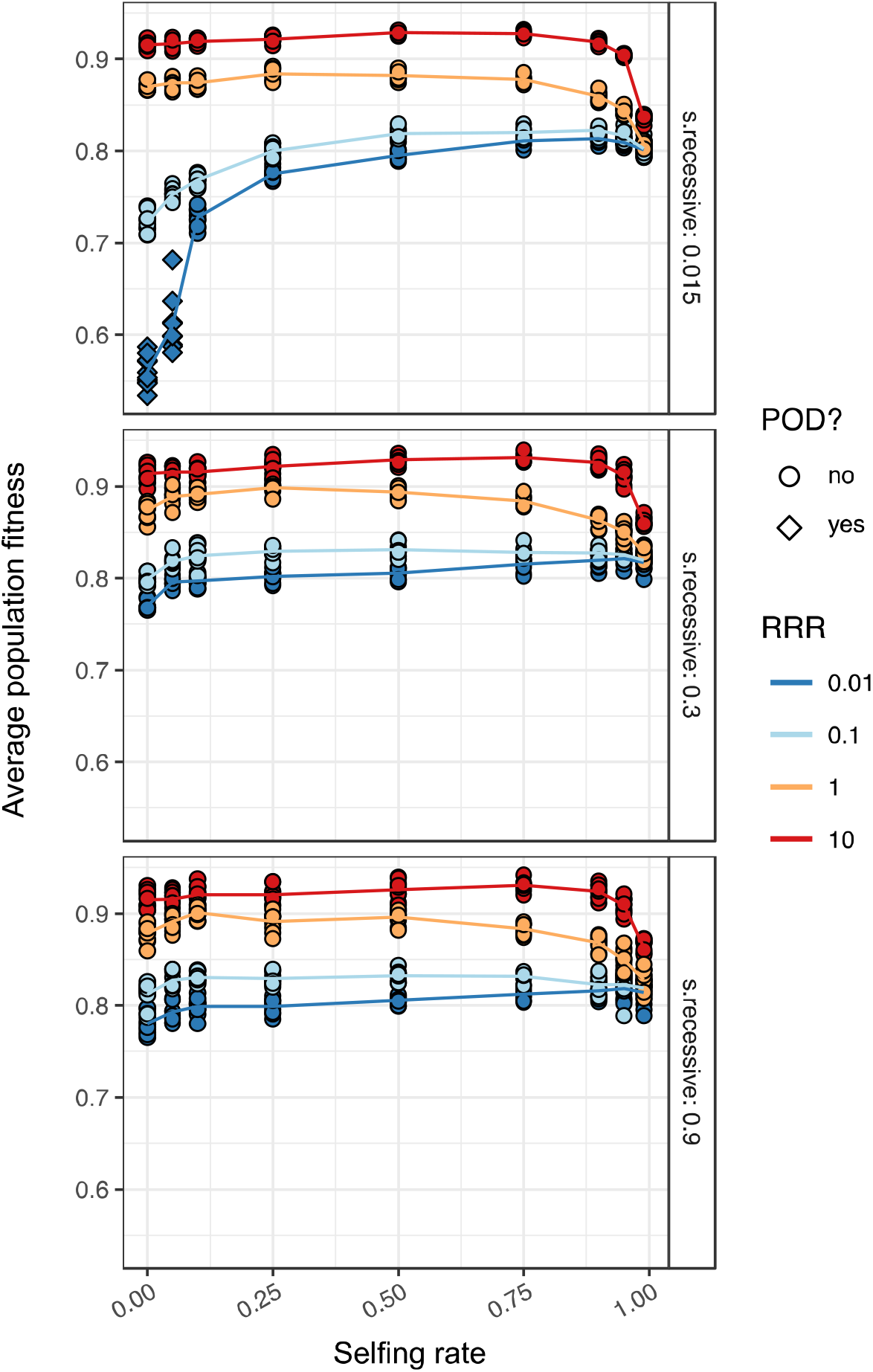
The relationship between mean population fitness and the selfing rate varies as a function of the recombination rate and the presence of pseudo-overdominance (POD). *U*_*del*_ = 0.04, *h*_*recessive*_= 0. The three facets represent different fitness effects of the recessive mutations. At this *U*_*del*_, POD only occurs when the recessive mutations are relatively mild (*s*_*recessive*_= 0.015).

## DISCUSSION

The evolution of self-fertilization is common, occurring in many animals, fungi, plants, protozoa, and algae (Jarne and Auld 2006; Hanschen *et al*. 2018). This transition provides the opportunity to test if/how numerous genomic features associated with the transition to selfing affect the efficacy of selection against deleterious mutations. The lack of consistent empirical evidence for reduced efficacy of selection in selfers (Haudry *et al*. 2008; Escobar *et al*. 2010; Slotte *et al*. 2010, 2013; Qiu *et al*. 2011; Hazzouri *et al*. 2012; Ness *et al*. 2012; Gioti *et al*. 2013; Brandvain *et al*. 2014) is often attributed to factors not directly related to mating system and/or the recency of most selfing lineages (Haudry *et al*. 2008; Glémin and Galtier 2012). These explanations may be true. However, our results highlight a limitation of narrowly focusing on one hypothesized consequence of selfing, as self-fertilization has numerous genomic consequences with different predicted effects for the efficacy of selection. Specifically, we discover that selection is more effective in outcrossing than selfing populations when recombination rates are not too low and recessive deleterious mutations are rare, but when recombination rates are low and highly recessive deleterious mutations are common, selection is more effective in (partial) selfers than outcrossers.

### Effects of selfing rate on the efficacy of selection against deleterious mutations

By jointly simulating deleterious recessive and additive mutations across a broad slice of parameter space, we found that increases in selfing rate can have positive, neutral or negative effects on the accumulation of genetic load (see Figures 5, S13).

#### The unlinked recessive load in partially selfing populations

When recombination rates are exceptionally high, the fate of mutations in nearly all populations (except for near obligate selfers which experience Mueller’s ratchet) is independent of any linked deleterious mutations because recombination rapidly dissociates an allele from its background. With these high recombination rates, some threshold selfing rate is required to purge the genetic load when mutation rates are high and mutations are severe (as in Lande *et al*. 1994; Kelly 2007). The prevalence of recessive mutations at selfing rates below this threshold value exceeds predictions from single locus theory; partial selfing generates correlations in homozygosity at unlinked loci which interferes with the purging process when multi-locus inbreeding depression is nearly lethal (i.e. the load cannot be purged when all selfed offspring die).

We find that the efficacy of selection decreases as populations approach the critical selfing rate required to purge the load. That is, when recombination rates are high and highly deleterious recessive mutations are common the additive load increases with the selfing rate until populations can purge their load. Once the selfing rate exceeds this purging threshold the additive load begins decreasing with the selfing rate (red and orange lines in the two lower left panels of Figures S11C and S12C). This finding is consistent with Sachdeva (2019), which finds a greater increase in the additive load in partially selfing populations in the presence of highly damaging recessive mutations than in cases without these recessive mutations.

#### A little outcrossing goes a long way when recombination is common

At intermediate recombination rates, mildly deleterious mutations accumulate in predominant selfers but are removed by selection in outcrossers and mixed maters. The low effective recombination rate in predominant selfers causes selection against additive mutations at one site to limit the efficacy of selection against other additive mutations at linked sites. By contrast mutations in mixed mating and outcrossing populations rapidly recombine away from linked deleterious mutations and selection at one site does not impact the efficacy of selection at linked sites (as seen in Charlesworth *et al*. 1993a; Glémin 2007; Glémin and Ronfort 2013; Kamran-Disfani and Agrawal 2014). Therefore, the weak empirical evidence for a decrease in the efficacy of selection in partially selfing populations may be partially attributable to the paucity of near obligate selfing in nature (Goodwillie *et al*. 2005; Moeller *et al*. 2017).

As the recombination rate becomes more similar to the mutation rate, the efficacy of selection starts to decrease more continuously with the selfing rate. This is because, at these lower recombination rates, mixed maters, but not outcrossers, begin to experience increased selective interference and background selection (Glémin 2007; Glémin and Ronfort 2013; Kamran-Disfani and Agrawal 2014). Consistent with our results and others, Laenen et al. (2018) found elevated load in only highly selfing (∼0.9 selfing rate) populations of *Arabis alpina*; with no increase in the load in mixed-mating (∼0.8 selfing rate) as compared to outcrossing populations.

#### Selection against alleles linked to deleterious recessive mutations is more effective is partially selfing populations

The equilibrium frequency of haplotypes without a deleterious mutation, *f*_*0*_, determines the strength of background selection and selective interference among linked deleterious mutations (Charlesworth *et al*. 1993a). By removing recessive mutations when rare, partial selfing increases *f*_*0*_ and decreases the extent of background selection and selective interference. In contrast, the accumulation of many, rare segregating recessive mutations in outcrossers decreases *f*_*0*_, decreasing *N*_*e*_ and lowering the efficacy of selection against linked deleterious mutations. Thus, with low relative recombination, selection against additive, mildly deleterious mutations becomes more effective as the selfing rate increases.

#### The shift from purifying selection to pseudo-overdominance weakens the efficacy of selection

When deleterious recessive mutations arise more quickly than selection and recombination can maintain any “unloaded haplotypes”, pseudo-overdominant haplotypes arise. In contrast to standard background selection driven by purifying selection, pseudo-overdominance increases diversity at linked neutral sites (Gilbert *et al*. 2020). The frequency of pseudo-overdominance in nature is unknown; however, a recent genome scan (Becher *et al*. 2020) identified numerous genomic regions displaying signatures of associative overdominance (which can be caused by pseudo-overdominance) in flies and humans, and a recent review (Waller 2021) compiled numerous lines of evidence suggesting that pseudo-overdominance is common in plants.

Although pseudo-overdominance increases diversity at linked neutral sites (Ohta and Kimura 1970; Gilbert *et al*. 2020), we find that it substantially increases the burden of deleterious mutations (see Figures 4 and 5). We propose that by sub-structuring a population into complementary haplotypes in repulsion, pseudo-overdominance effectively decreases the N_e_ that affects the efficacy of selection, as is generally predicted in subdivided populations (Whitlock 2003). Because the pseudo-overdominant haplotypes form in low recombination regions, there is effectively no ‘migration’ of alleles between haplotypes (Charlesworth *et al*. 2003; Charlesworth 2006). The consequence is that the *N*_*e*_ that determines the efficacy of selection against mildly deleterious additive mutations is thus a function of the number of genomes *within* a haplotype class.

Once pseudo-overdominant haplotypes emerge, additional recessive mutations are sheltered from selection and continue to accumulate, as is known for other cases of heterozygote advantage (Mather and de Winton 1941; Glémin *et al*. 2001; van Oosterhout 2009; Jay *et al*. 2021). This sheltered load can reinforce pseudo-overdominance, because genomic regions which are rarely homozygous are free to accumulate additional recessive variants (Llaurens *et al*. 2017), which further increases the strength of selection against individuals homozygous in these regions. Such a pattern has been shown for certain types of polymorphic inversions (Berdan *et al*. 2021).

#### Selfing prevents the shift from background selection to pseudo-overdominance

The analytical theory derived here qualitatively matches results from our individual-based simulations and shows that (1) by purging the recessive load, partial selfing prevents a shift from purifying selection to pseudo-overdominance, and (2) that recombination amplifies the effects of partial selfing on preventing the transition to pseudo-overdominance. Overall, we find a sharp decrease in the parameters allowing for pseudo-overdominance in partially selfing populations (Figure 2E). At a given partial selfing rate (i.e., selfing rate < 0.5), pseudo-overdominance becomes more likely when *U*_*del*_ is high and *s*_*recessive*_ is low, as these are parameter combinations where it is harder to purge the recessive load (Wang *et al*. 1999). As populations experience a greater influx of deleterious recessive mutations, a higher selfing rate is needed to purge the recessive load before it becomes structured into complementary, pseudo-overdominant haplotypes.

### Caveats and future directions

#### The joint distribution of dominance and fitness effects

We simulated populations with four equally probable mutations – three types of mildly deleterious additive mutation types, and one strongly deleterious (partially) recessive mutation type. This mutational model is obviously wrong. In reality mutations take selective and dominance coefficients from a two-dimensional density function. The best methods to infer the distribution of fitness effects from polymorphism data (Keightley and Eyre-Walker 2007) provide only crude estimates of this distribution. However, two of the best studies on the topic show that more recessive mutations are more deleterious (Agrawal and Whitlock 2011; Huber *et al*. 2018).

Our chosen parameters, consisting of recessive mutations with selection coefficients much larger than 4*Ns*, and additive mutations with selection coefficients closer to 4*Ns*, capture the spirit of this result. Still, because highly damaging mutations are unlikely to be fully recessive (Crow 1993), it is worth noting that most qualitative results found with complete recessivity are also found when *h =* 0.1. In fact, pseudo-overdominance can emerge when *h =* 0.1 (Figures S2B and S6A).

#### Demographic history

Factors affecting the efficacy of selection other than the automatic genomic consequences investigated here often change with the mating system. For example, selfing is often associated with colonization of and rapid expansion in islands, disturbed, or other marginal habitats (Baker 1955), further decreasing the efficacy of selection in selfers. Therefore, selfers may suffer a more severe expansion load (Peischl *et al*. 2013, 2015) than outcrossers. However, demographic changes such as population expansion and contraction have more influence on recessive than additive mutations, making their effects on (partially) selfing populations likely limited (Kirkpatrick and Jarne 2000; Balick et al. 2015; Peischl et al. 2015). Nonetheless, an integration of both the genetic and demographic consequences of the mating system would better predict differences in the genetic load associated with the mating system.

#### Realistic genomic architecture

We assumed that recombination and mutation rates did not vary across the genome. In reality, however, recombination and deleterious mutation rates vary across the genome (Gaut *et al*. 2007; McVicker *et al*. 2009; Slotte 2014) and can positively (e.g., *Mimulus guttatus* (Aeschbacher *et al*. 2017), maize (Anderson *et al*. 2006), rice (International Rice Genome Sequencing Project, Takuji Sasaki 2005), wheat (Dvorak *et al*. 2004), *A. thaliana* (Wright *et al*. 2003; Giraut *et al*. 2011), and *Populus* species (Wang *et al*. 2016; Apuli *et al*. 2020) or negatively (e.g., *Caenorhabditis* (Barnes *et al*. 1995), and *Mimulus aurantiacus* (Stankowski *et al*. 2019)) covary. Future work could address how the results described here translate into differences in the load across genomes as a function of the association between gene density and recombination rates.

## Conclusions

We highlight the multifaceted pathways in which (partial) selfing affects the efficacy of selection against deleterious mutations. The effect of mating system on the efficacy of selection is primarily driven by interactions between dominance coefficients and the rates of selfing, recombination and mutation. We find that the joint consideration of mutations with either recessive or additive effects on fitness importantly changes the relationship between genetic load and selfing rate, as the strength of linked selection driven by mutations of either dominance class varies with selfing rate. In particular, we find that a shift from classic purigying selection to pseudo-overdominance in primarily outcrossing populations drastically reduces the efficacy of selection against mildly deleterious additive mutations and that partial selfing prevents a shift to pseudo-overdominance, resulting in another way by which genetic load decreases with selfing rate.

