## Supplementary table and figures for "Genetic load may increase or decrease with selfing depending upon the recombination environment"

**Table S1**: Translation of the genome-wide deleterious mutation rate (U_del_) and the relative recombination rate (RRR) parameters to other measures of the mutation rate (per-base pair) and recombination rate (per-base pair, and cM/Mb).

| **U_del_** | **RRR** | **mutation rate** |  | **recombination rate** | |
| --- | --- | --- | --- | --- | --- |
|  |  | per-base-pair |  | per-base-pair | cm/Mb |
| 0.04 | 0.01 | 8.89E-10 |  | 8.89E-12 | 0.0089 |
| 0.04 | 0.1 | 8.89E-10 |  | 8.89E-11 | 0.0889 |
| 0.04 | 1 | 8.89E-10 |  | 8.89E-10 | 0.8889 |
| 0.04 | 10 | 8.89E-10 |  | 8.89E-09 | 8.8889 |
| 0.16 | 0.01 | 3.56E-09 |  | 3.56E-11 | 0.036 |
| 0.16 | 0.1 | 3.56E-09 |  | 3.56E-10 | 0.356 |
| 0.16 | 1 | 3.56E-09 |  | 3.56E-09 | 3.556 |
| 0.16 | 10 | 3.56E-09 |  | 3.56E-08 | 35.556 |
| 0.48 | 0.01 | 1.07E-08 |  | 1.07E-10 | 0.11 |
| 0.48 | 0.1 | 1.07E-08 |  | 1.07E-09 | 1.07 |
| 0.48 | 1 | 1.07E-08 |  | 1.07E-08 | 10.67 |
| 0.48 | 10 | 1.07E-08 |  | 1.07E-07 | 106.67 |

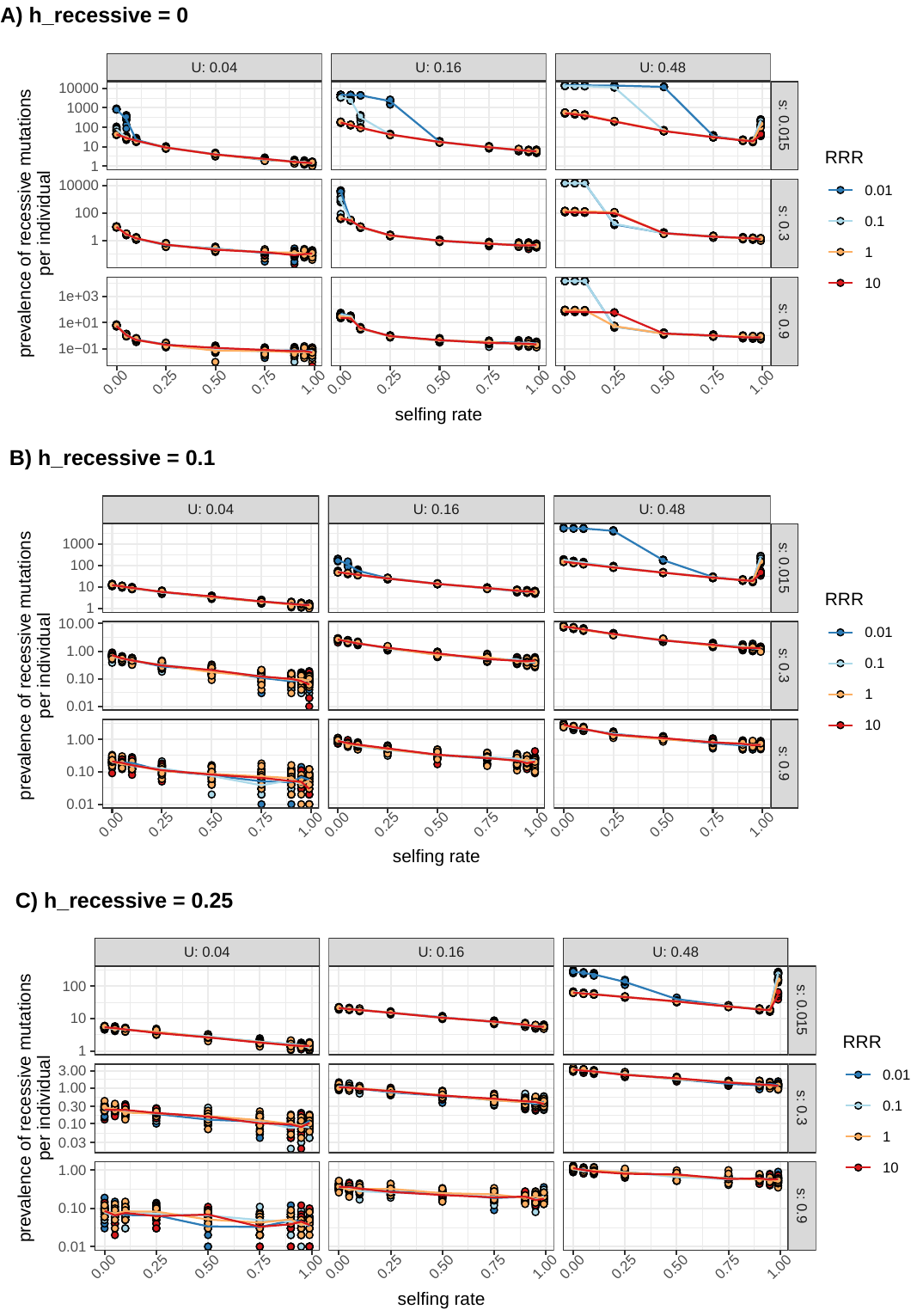

**Figure S1**: Purging dynamics of recessive mutations as a function of selfing rate in simulations with high (warm colors) vs low (cool colors) relative recombination rates (RRR) highlights the qualitative spike in recessive load due to pseudo-overdominance. Points indicate the average prevalence of recessive mutations per individual in a given simulation replicate, and lines connect the mean value among simulation replicates. A) h_recessive_=0, B) h_recessive_=0.1, C) h_recessive_=0.25. Within each panel, columns indicate different genome-wide mutation rates (U_del_) and rows indicate the deleterious fitness effects of recessive mutations (s_recessive_). Note log10 scale on the y-axis.

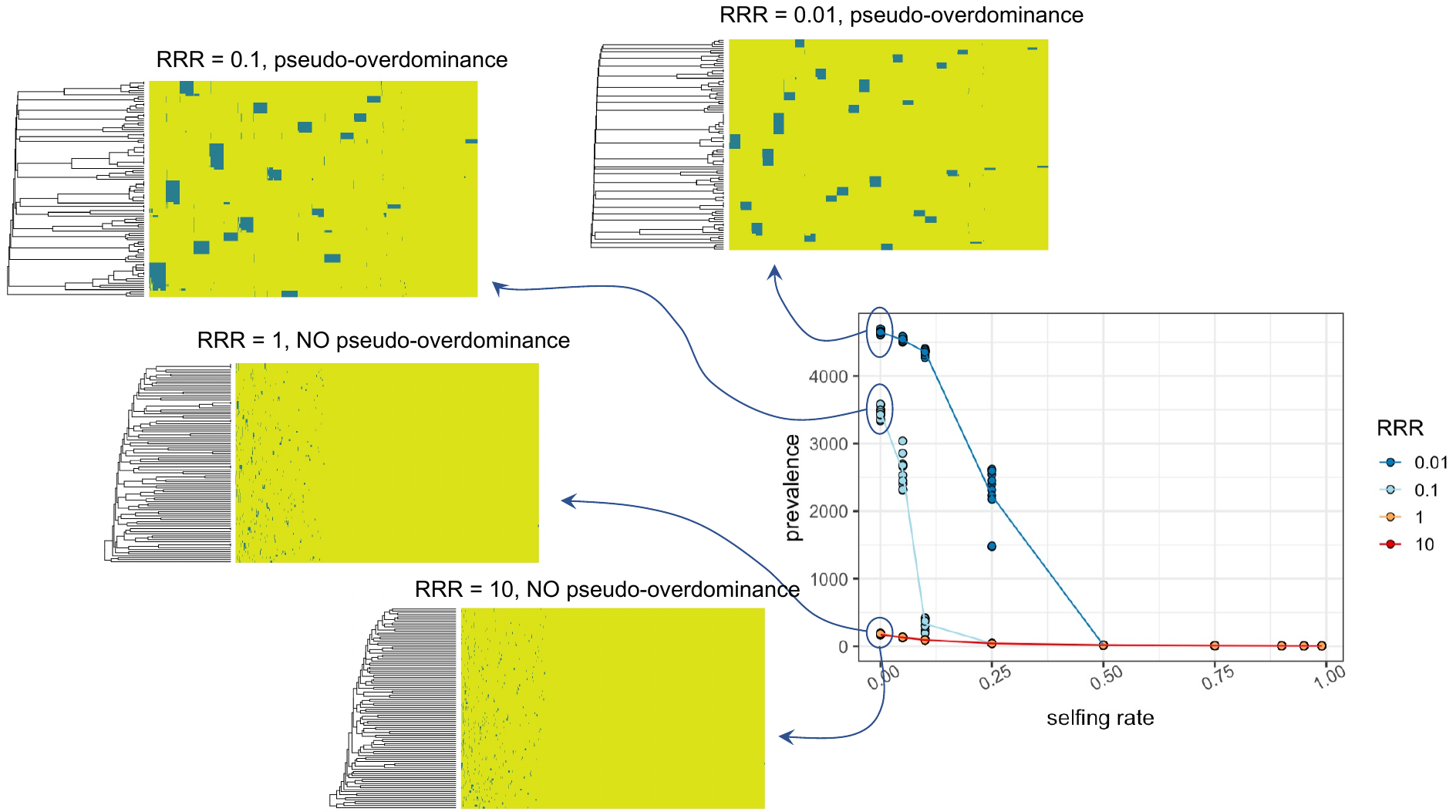

**Figure S2:** Clustering analyses confirm that pseudo-overdominance occurs at low recombination rates and leads to haplotype structuring within populations. The graph on the right shows the prevalence of recessive deleterious (s_recessive_=0.015) mutations as a function of selfing rate. At a selfing rate of zero, for example, we called pseudo-overdominance at relative recombination rates (RRR) of 0.01 and 0.1, but not at 1 or 10, using indirect evidence from the prevalence of recessive mutations, neutral diversity (pi) and the allele frequency spectrum. To confirm that, indeed, pseudo-overdominance is occurring, we visualized haplotype structure in simulations at a selfing rate of zero across all RRRs. Heatmaps show the presence (blue) or absence (green) of all recessive mutations in a simulation (columns) for 100 sampled haplotypes (rows). Dendrograms on the left show the relatedness among haplotypes, and demonstrate strong clustering of haplotypes in simulations with pseudo-overdominance. Columns are also clustered and thus do not represent genomic position.

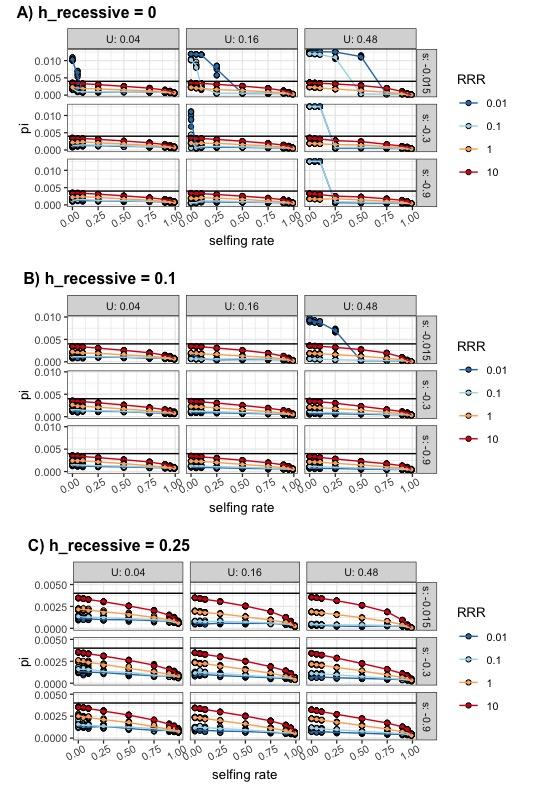

**Figure S3:** Neutral diversity (π) spikes in simulations with pseudo-overdominance, and otherwise decreases with selfing rate and the relative recombination rate (RRR). Black horizontal lines indicate the neutral expectation of π (= 4Nμ). A) h_recessive_=0, B) h_recessive_=0.1, C) h_recessive_=0.25. Within each panel, columns indicate different genome-wide mutation rates (U_del_) and rows indicate deleterious fitness effects of recessive mutations (s_recessive_).

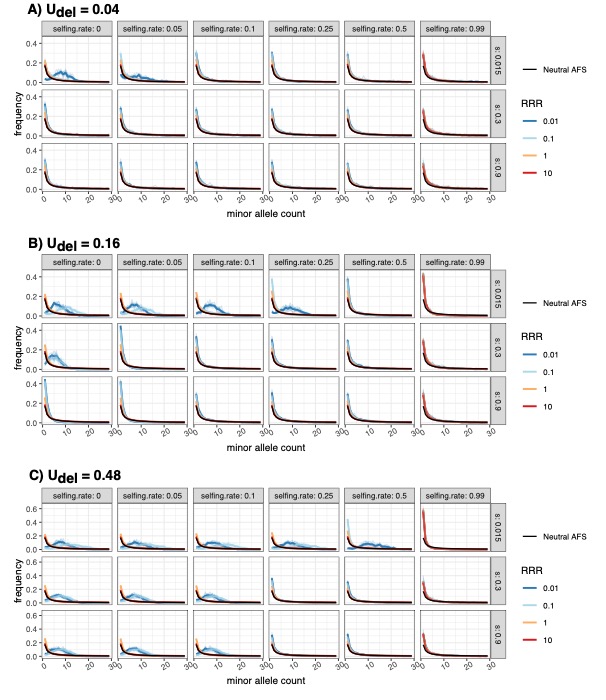

**Figure S4:** A snapshot of allele frequency spectra (AFS) in simulations with fully recessive (h_recessive_=0) mutations. Within each panel (A-C; varying mutation rates), each graph is the AFS for a given selfing rate (column) and fitness effect of recessive mutations (s_recessive_, row). Black lines indicate expected neutral AFS. We show the frequency of alleles that occur up to 30 times in a sample of 200 genomes.

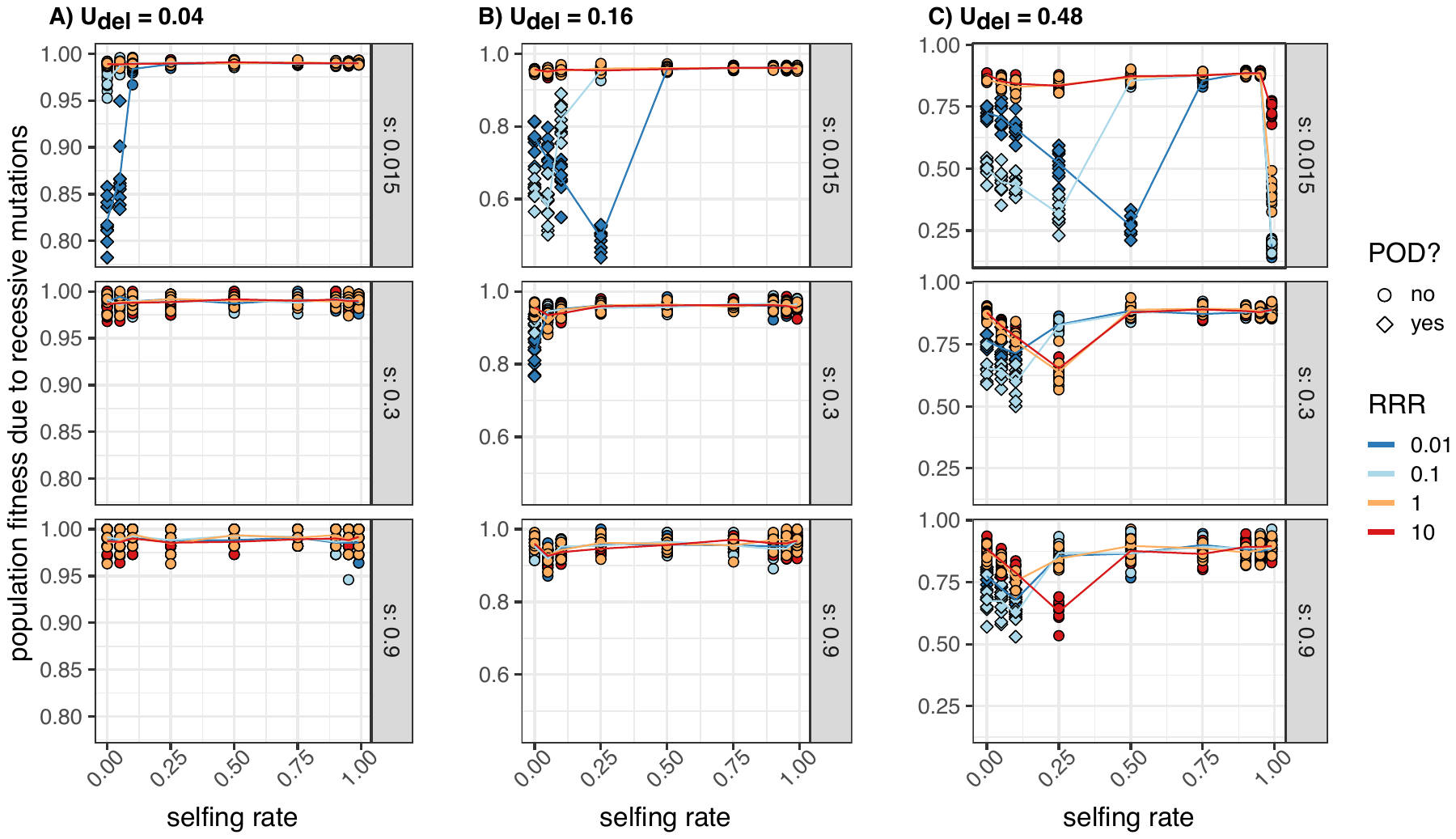

**Figure S5:** Average population fitness due to recessive (*h_recessive_* = 0) mutations across the three genome-wide deleterious mutation rates (A-C) and the selection coefficient of the recessive mutations (s_recessive_, rows). Drops in fitness in low recombination environments (RRR <1) are due to the presence of pseudo-overdominance.

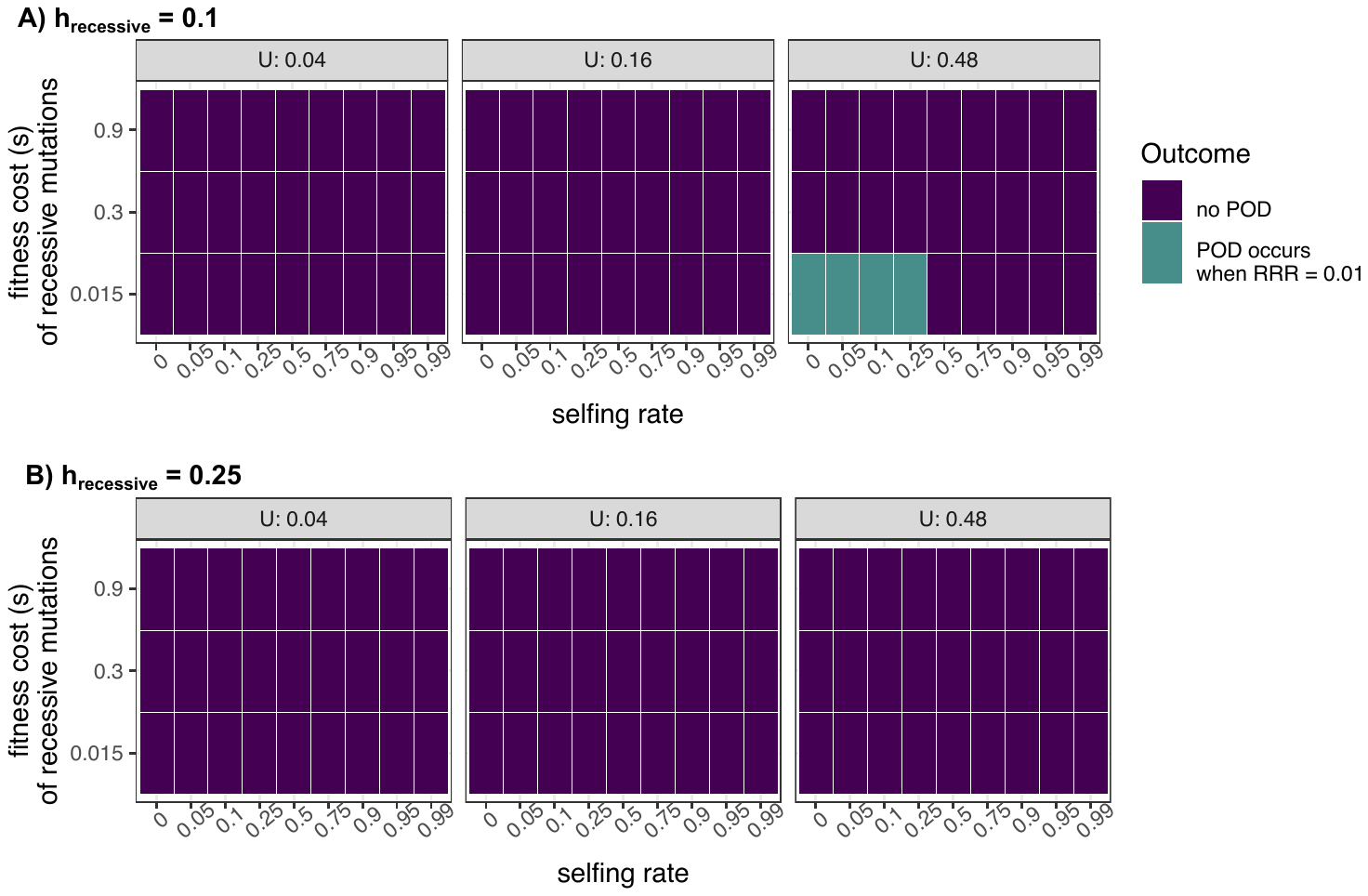

**Figure S6:** Outcome plot indicating when pseudo-overdominance (POD) occurs across parameter space for partially recessive mutations. (A) Dominance of recessive mutations (h_recessive_) = 0.1. (B) Dominance of recessive mutations (h_recessive_) = 0.25.

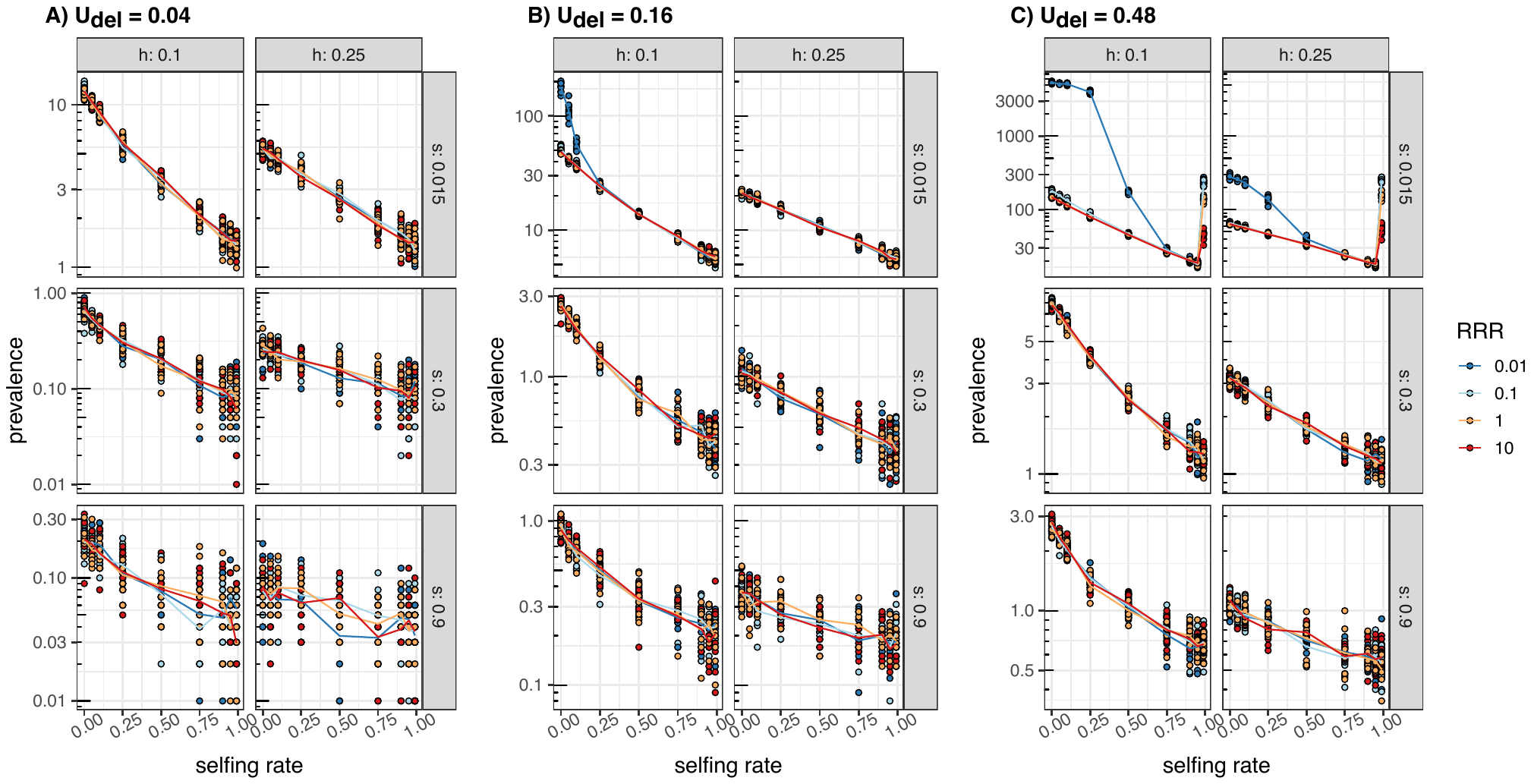

**Figure S7**: Prevalence of recessive mutations (average prevalence per individual) as a function of selfing rate when mutations are partially recessive (*h_recessive_* = 0.1 or *h_recessive_* = 0.25) at the three genome-wide deleterious mutation rates (A-C). Rows within panels show prevalence for different fitness effects of the recessive mutations (s_recessive_).

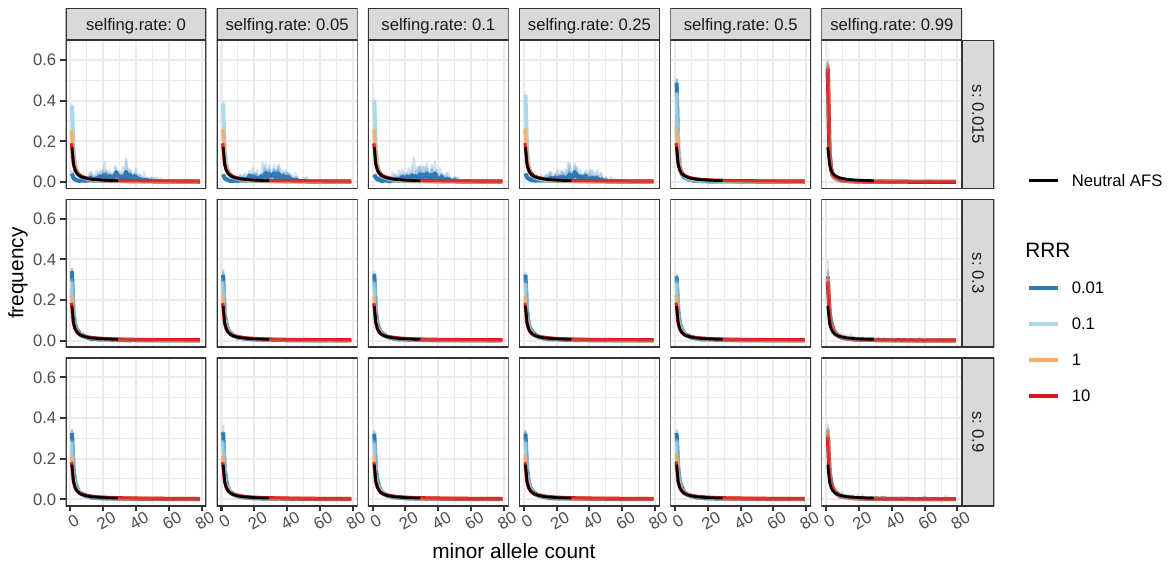

**Figure S8:** Allele frequency spectra when *h_recessive_=* 0.1 and U_del_ = 0.48. We show only U_del_ = 0.48, because at this h_recessive_, pseudo-overdominance only occurs at U_del_ = 0.48. The shift to pseudo-overdominance is evidenced by the shift to more intermediate allele frequencies at the lowest relative recombination rate (RRR) when the selfing rate (columns) is less than 0.5 and the fitness effect of recessive mutations (rows) is 0.015. Black lines indicate expected neutral AFS. We show the frequency of alleles that occur up to 30 times in a sample of 200 genomes.

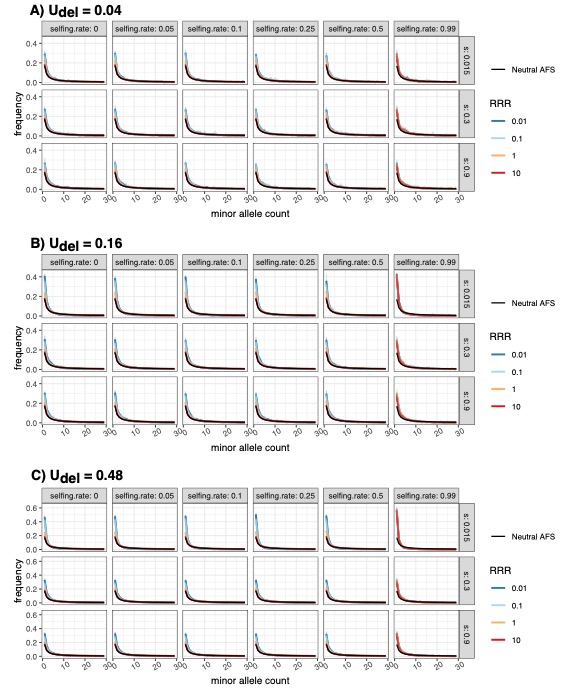

**Figure S9:** A snapshot of allele frequency spectra (AFS) in simulations with partially recessive (*h_recessive_*= 0.25) mutations. Within each panel (A-C; varying mutation rates), each graph is the AFS for a given selfing rate (column) and fitness effect of recessive mutations (s_recessive_, row). Black lines indicate expected neutral AFS. We show the frequency of alleles that occur up to 30 times in a sample of 200 genomes.

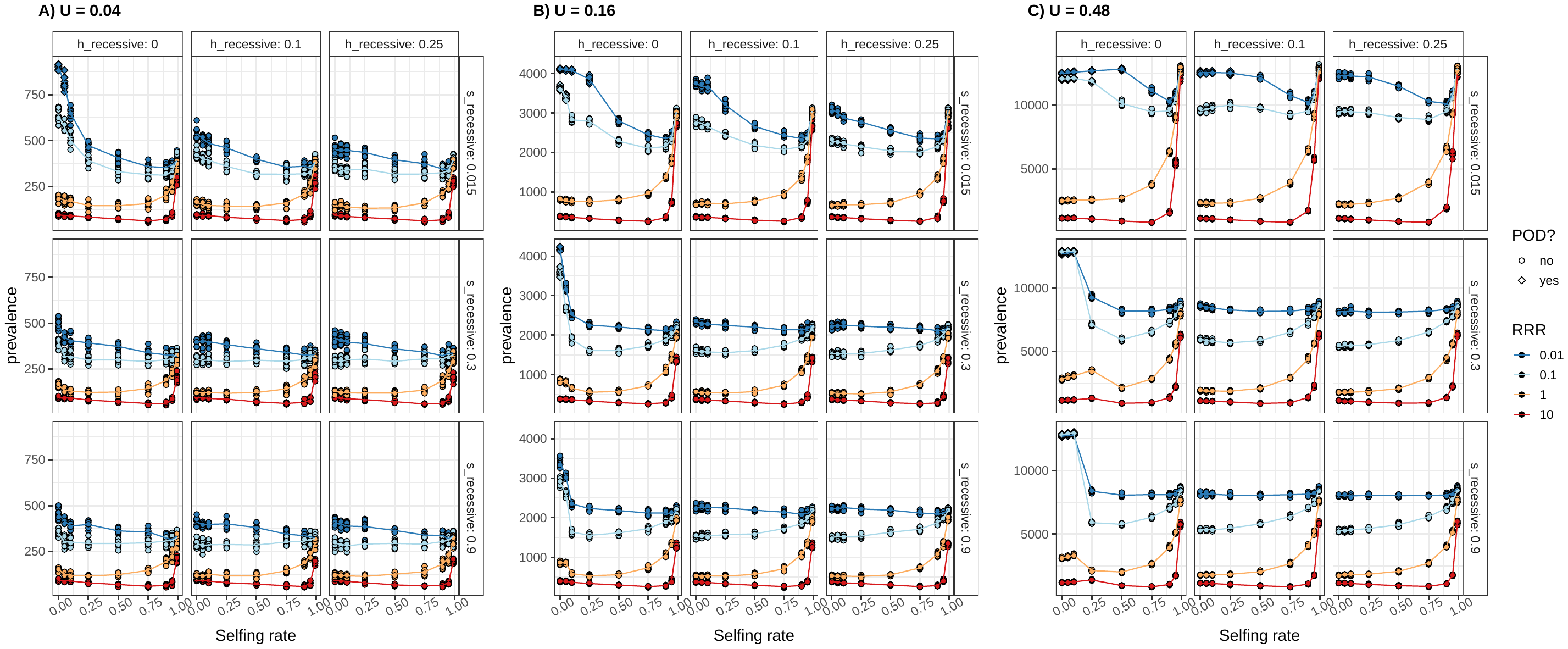

**Figure S10:** Prevalence of additive mutations characterized by s = -0.0005 across the three genome-wide deleterious mutation rates (A-C). Columns within each panel characterize the dominance of recessive mutations present in the simulation. The presence of pseudo-overdominance affects the accumulation of additive load, as seen by the differences in prevalence between simulations with fully and partially recessive mutations, particularly in low recombination simulations. Rows within each panel characterize the fitness effect of recessive mutations.

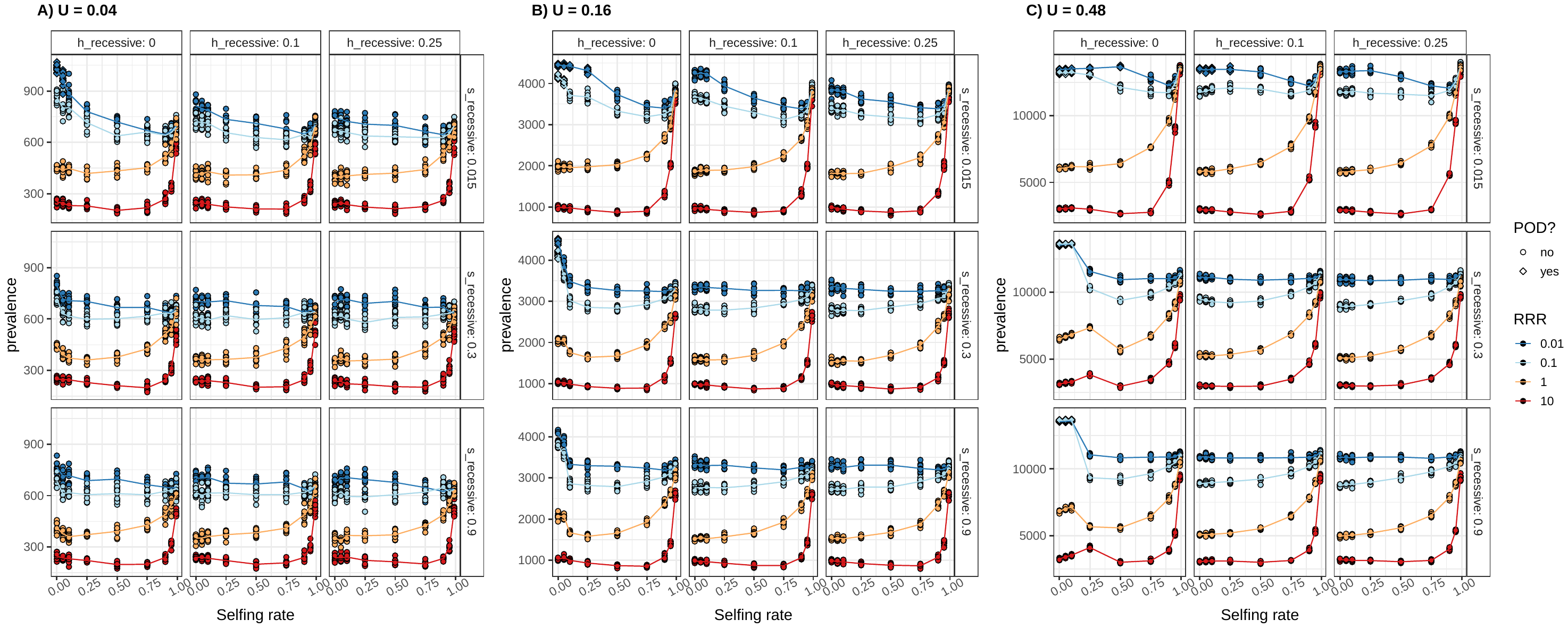

**Figure S11:** Prevalence of additive mutations characterized by s = -0.00025 across the three genome-wide deleterious mutation rates (A-C). Columns within each panel characterize the dominance of recessive mutations present in the simulation. The presence of POD affects the accumulation of additive load, as seen by the differences in prevalence between simulations with fully and partially recessive mutations, particularly in low recombination simulations. Rows within each panel characterize the fitness effect of recessive mutations.

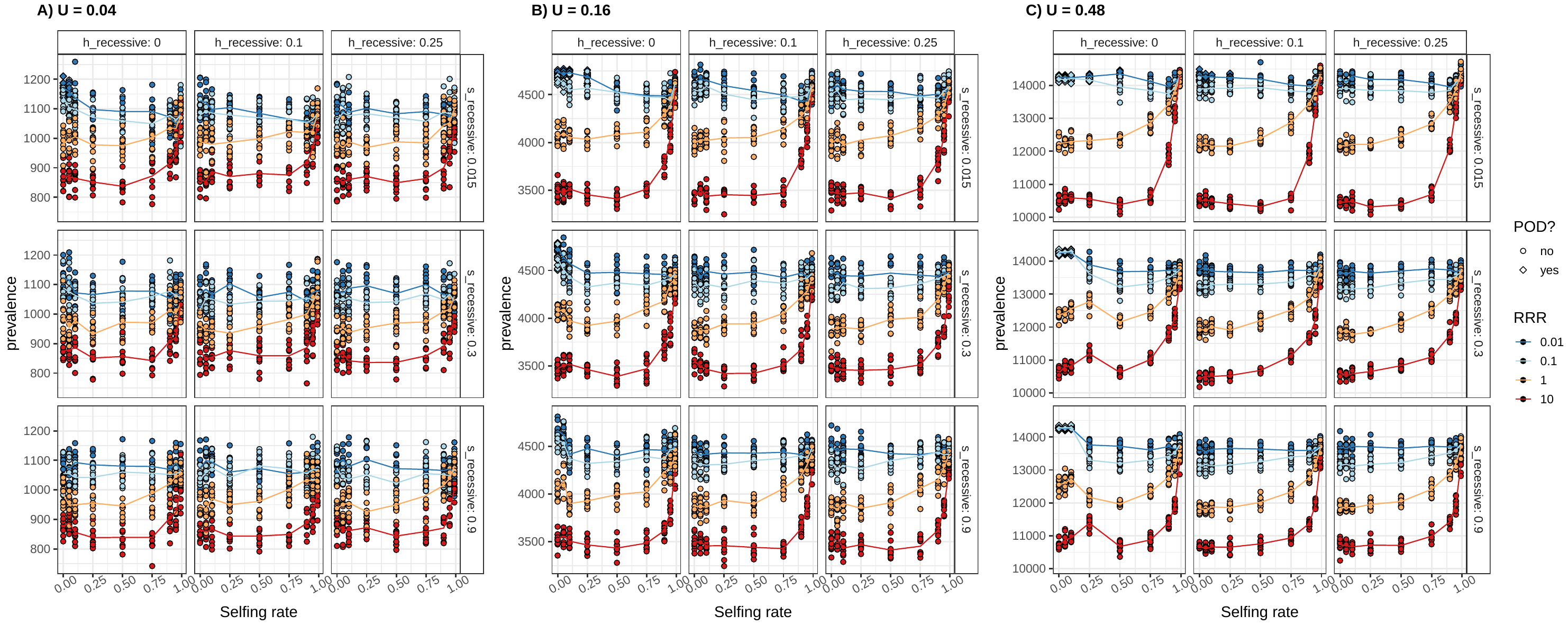

**Figure S12:** Prevalence of additive mutations characterized by s = 0.00005 across the three genome-wide deleterious mutation rates (A-C). Columns within each panel characterize the dominance of recessive mutations present in the simulation. The presence of POD affects the accumulation of additive load, as seen by the differences in prevalence between simulations with fully and partially recessive mutations, particularly in low recombination simulations. Rows within each panel characterize the fitness effect of recessive mutations.

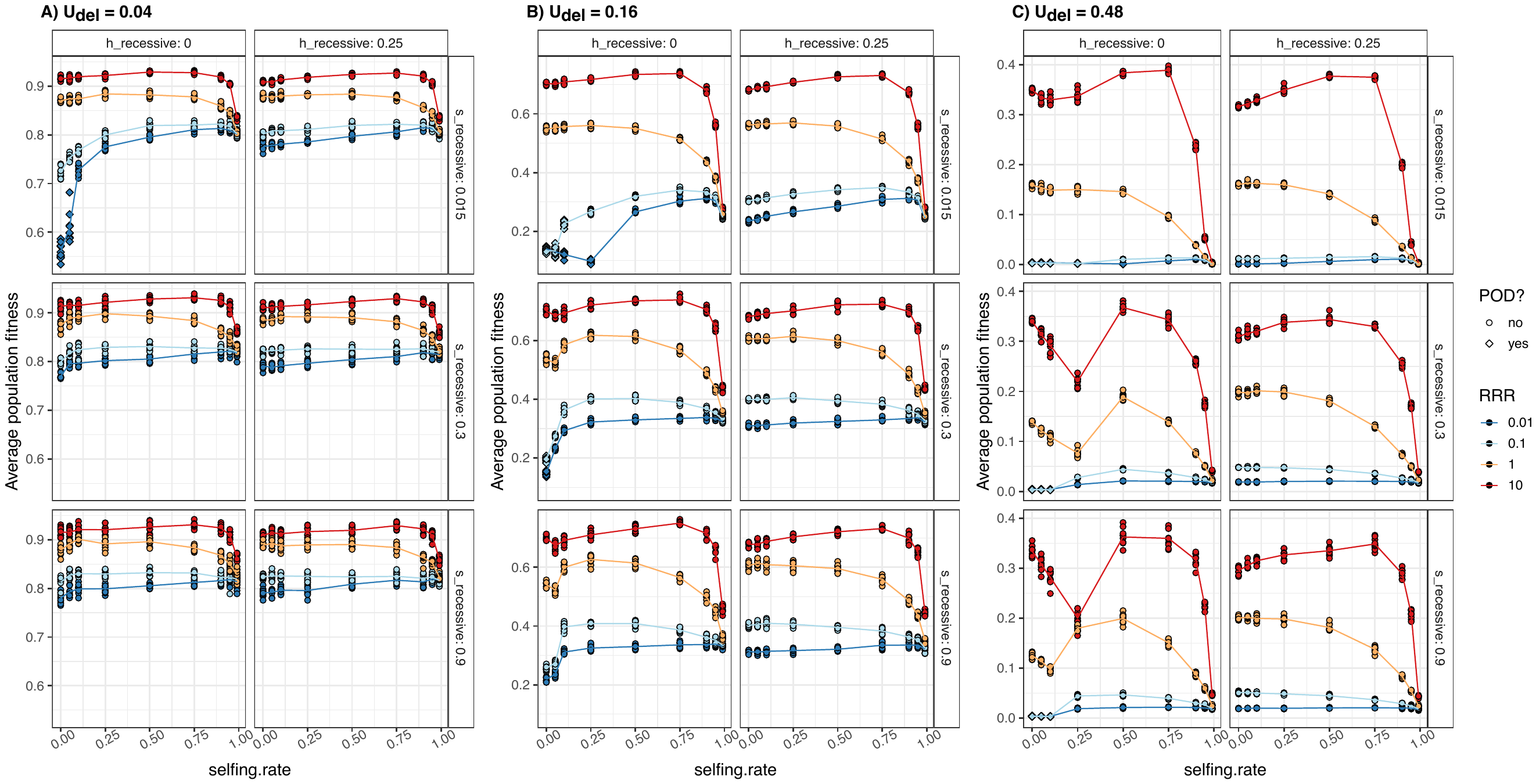

**Figure S13:** Mean population fitness across the three genome-wide mutation rates (A-C). Columns within each panel characterize the dominance of recessive mutations present in the simulation. The presence of POD affects population fitness, as seen by the differences in prevalence between simulations with fully and partially recessive mutations, particularly in low recombination simulations. Rows within each panel characterize the fitness effect of recessive mutations.
