## Appendix for "Genetic load may increase or decrease with selfing depending upon the recombination environment"

### ***A model for mutation load in partially selfing populations***

#### **Genotype Frequencies with Selfing**

We extend the multi-locus model from Gilbert et al. (2020) to include selfing. We consider $n$ biallelic loci. We denote wild-type and derived alleles alleles at locus $i=1.,...,n$ by $a_{i}$ and $A_{i}$, respectively. Derived alleles are deleterious and fully recessive with fitness $1-s$ when homozygous and $1$ otherwise. There is no epistasis and we assume multiplicative effects across loci.

For fully linked loci (that is, there is no recombination) the model is equivalent to a single locus model with $2^{n}$ alleles where each haplotype corresponds to a single allele. As in Gilbert et al. (2020), we want to derive the frequency of the 0-mutation haplotpye $p_{0}$ at mutation selection balance. When this frequency approaches 0, we expect a transition to pseudo-overdominance (POD). We assume that haplotypes with more than one deleterious mutations are vanishingly rare at equilibrium. This has the important consequence that no genotype can be derived homozygous at more than one locus, which simplifies calcualtions considerably. Let $p_{i}$, $i=1,...,n$ be the frequency of the haplotype carrying a derived allele at locus $i$, and $p_{0}$ the frequency of the haplotype without any deleterious mutations.

With these haplotypes, we only need to follow four different multi genotypic categories (Caballero and Hill 1992):

1. A genotype without any deleterious mutations. This genotype is found at frequency $p_{00}=F\sum_{i=0}^{n} p_{0}p_{i}+(1-F)p_{0}^{2}$.
2. A genotype heterozygous at one locus. This genotype is found at frequency $p_{01}^{(1)}=2(1-F)\sum_{i=1}^{n} p_{0}p_{i}$.
3. A genotype heterozygous at two loci. This genotype is found at frequency $p_{01}^{(2)}=(1-F)\sum_{i=1}^{n} \left( \sum_{j=i+1}^{n} 2p_{i}p_{j} \right)$. And
4. Genotypes that are derived homoyzgous at one locus $p_{11}=\sum_{i=1}^{n} \left( F\sum_{j=0}^{n} p_{i}p_{j}+(1-F)p_{i}^{2} \right)$.

Fitness of these genotype-classes is given by $w_{00}=w_{01}^{(1)}=w_{01}^{(2)}=1$ and $w_{11}=1-s$. Mean fitness is given by $\overline{w}=1-sp_{11}$. Assuming continuous time the change in genotype frequencies after selection are then given by $dp_{g}/d_{t}=p_{g}(w_{g}-\overline{w})$, $g\in\{00,{}_{01}^{(1)},{}_{01}^{(2)},11\}$. The parameter $F$ is equivalent to $\alpha/(2-\alpha)$ where $\alpha$ is the selfing rate (Caballero and Hill, 1992).

#### **Evolutionary Dynamics of** $\boldsymbol{p}_{\boldsymbol{0}}$ **with recombination**

At equilibrium all haplotypes that carry a deleterious allele must have the same frequency, which we denote as $p_{1}$. That is, from now on we simply use $p_{1}$ to refer to the frequency of any haplotype carrying exactly one derived allele. At equilibrium $p_{1}=(1-p_{0})/n$ such that haplotype frequencies sum up to 1. Since we are only interested in calculating haplotype frequencies at mutation-selection equilibrium we henceforth assume that $p_{1}=(1-p_{0})/n$. Solving

$$\frac{dp_{0}}{d_{t}}=0$$

then yields the equilibrium frequency of the zero-mutation haplotype $\hat{p}_{0}$.

We next assume that loci are equidistantly distributed over a region of length $r$ cM, such that the recombination rate between adjacent loci is $r/(n-1)$. We assume that recombination is sufficiently rare such that we can ignore double recombination events. Since we ignore haplotypes with more than one deleterious allele, the main effect of recombination is to restore haplotypes without any deleterious alleles from genotypes heterozygous at two loci. Analogous to Gilbert et al. (2020) the equation for the equilibrium haplotype frequency then becomes:

$$\frac{dp_{0}}{d_{t}}=p_{00}(w_{00}-\overline{w})-nup_{0}+\frac{1}{2}p_{01}(w_{01}-\overline{w})+\sum_{i=1}^{n} \left[ \sum_{j=i}^{n} \left( (1-F)\frac{r(j-i)}{(n-1)}p_{i}p_{j}+F\frac{r(j-i)}{(n-1)}\frac{p_{11}}{n} \right) \right]=0.$$

Here, $u$ is the per locus mutation rate and $\frac{r(j-i)}{(n-1)}$ is the recombination rate between loci $i$ and $j$.

#### **Special cases**

##### **Single Locus, No Selfing**

For a single locus without selfing ($n=1,F=0$) we get

$$\frac{dp_{0}}{d_{t}}=p_{0}\left( (p_{0}-1)^{2}s-u \right)$$

and $\hat{p}=1-\sqrt{u/s}$, recovering the well-known result for mutation-selection balance of recessive alleles. For $n$ loci, $\hat{p}=1-n\sqrt{u/s}$, in agreement with previous results (Gilbert et al., 2020).

##### **No Recombination**

For partially or fully selfing organisms without recombination we get

$$\frac{dp_{0}}{d_{t}}=-\frac{p_{0}\left[ (p_{0}-1)s(F(n+p_{0}-1)-p_{0}+1)+n^{2}u \right]}{n}$$

and

$$\hat{p}_{0}=1-\frac{n\left( \frac{\sqrt{F^{2}s-4Fu+4u}}{\sqrt{s}}-F \right)}{2(1-F)}.$$

For $n=1$ this gives

$$1-\frac{\frac{\sqrt{F^{2}s-4Fu+4u}}{\sqrt{s}}-F}{2(1-F)},$$

recovering the results from Roze and Rousset (2004). In particular, the frequency of the deleterious allele at equilibrium is

$$\hat{p}_{1}=(1-\hat{p}_{0})=\frac{\frac{\sqrt{F^{2}s-4Fu+4u}}{\sqrt{s}}-F}{2(1-F)}.$$

Using the fact that $F=\frac{\alpha}{2-\alpha}$ we get

$$\hat{p}_{1}=\frac{\alpha\sqrt{s}+(\alpha-2)\sqrt{\frac{\alpha^{2}s+8(\alpha-2)(\alpha-1)u}{(\alpha-2)^{2}}}}{4(\alpha-1)\sqrt{s}}, (A1)$$

which is equivalent to the formula given in Table 1 of Roze and Rousset (2004). Furthermore, the mutation load is then simply $u$ for completely recessive mutations. Figure A1 shows the frequency of the deleterious allele at mutation selection balance.

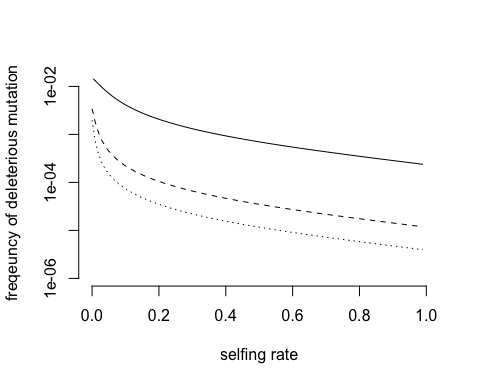

**Figure A1.** Frequency of deleterious mutation at mutation selection balance. Parameter values are u = 3.5 * 10^-6^ and s = 0.015, 0.3, 0.9 (from top to bottom).

##### **Multiple Loci, Full Selfing**

For fully selfing populations we get:

$$\hat{p}_{0}=1-n\frac{u}{s}, (A2)$$

and hence find that mutation load is independent of recombination. This is sensible as recombination can only act on heterozygotes, which are absent at equilibrium in fully selfing populations.

#### **Recombination and selfing**

Direct analytical progress seems unfeasible and we hence resort to approximation. We assume that evolutionary forces are weak, that is $s,u,$ and $r<<1$. This also means that $n$ cannot be too large because with weak recombination multi-locus genotypes might experience selection coefficients of up to $ns$. Furthermore, we assume that recombination rate $r$ and mutation rate $u$ are small relative to the strength of selection and selfing. Our approximation should be reliable if selection is weak, but larger than $1/N$ where $N$ is the population size. Furthermore, recombination and per locus mutation rates should be weak relative to selection. The exact conditions for when our approximation will become unreliable are difficult to derive however. Roughly speaking $r,u<1/N<<s<<1$ is assumed throughout this section. Our approximation may not be useful for predictions on a genomic scale, but can accurately describe the dynamics of smaller regions of low recombination and should thus be helpful in providing qualitative explanations.

##### **Weak recombination and selfing**

Using an approach similar to that of Gilbert et al. (2020) we find that

$$\hat{p}_{0}=1-\sqrt{\mu}n+\frac{n}{2}\left( F+2\sqrt{\mu}-\sqrt{F^{2}+4\mu} \right)+\frac{1}{12}\sqrt{\mu}(n+1)n\rho, (A3)$$

where we use the re-scaled parameters $\mu=m/s,\rho=r/s$. The solution is obtained by using the Ansatz $\hat{p}_{0}=1+C_{1}\sqrt{\mu}+C_{2}\rho+C_{3}\varphi$, where $\varphi=F/s$. Then we develop $\frac{dp_{0}}{d_{t}}$ in a Taylor Series around $\rho=0$ while keeping the ratios $\rho/\sqrt{\mu}$ and $\rho/\varphi$ constant. Finally, we solve for $C_{1}$, $C_{2}$ and $C_{3}$.

Figure A2 compares the analytic approximation with 2 locus simulations and shows that the approximation is very accurate in the case without recombination. With increasing recombination or selfing rates the fit between theory and simulations becomes increasingly worse, but the transition from the region where we do not expect POD ($P_{0}>0$) to the region where we expect a transition to POD ($P_{0}=0$) is accurately predicted.

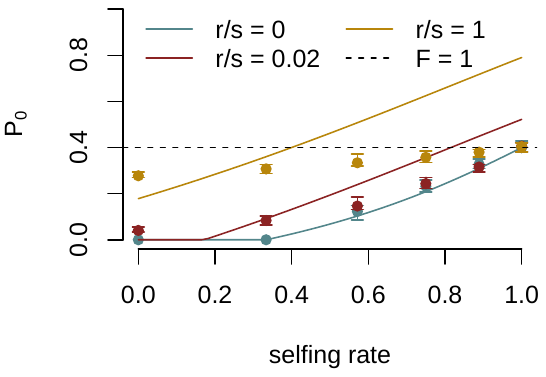

**Figure A2.** Frequency of the unloaded haplotype as a function of the selfing rate. Parameter values are n = 2 loci, u/s = 0.3, s = 0.01, N = 5000 diploid individuals and recombination rate r as specified in the legend. Points show the median and error bars the 95 % CI from 50 independent simulation replicates per parameter combination. Solid lines show the analytical approximation equation (A3) and dashed line the analytical expression for F = 1, equation (A2).

We can now use this formula to find the region in the parameter space for which we expect POD to occur. This is when $\hat{p}_{0}$ approaches 0. Figure A3 illustrates this region.

We note that it is difficult to directly compare our theoretical results to genome-wide simulations. First, our approximation will generally not be valid for small selection coefficients and/or large mutation rates as we have to assume weak evolutionary forces to derive the mutation-selection balance equilibrium. Second, our model assumes that mutation rates are small relative to selection coefficients at each locus, which might be violated in low recombination regions that accumulate numerous deleterious mutations with a large net fitness effect. Third, our model does not distinguish between loss of the zero-mutation haplotype and an actual transition to POD. While loss of the zero mutation-haplotype is necessary for POD to occur, it may also be the case that selection is so weak relative to mutation and recombination that there is subsequent fixation of deleterious mutations (Lynch *et al.* 1995), which would not be classified as POD in the simulations. In the main text we nevertheless show a qualitative comparison to individual based simulations and find that our model can accurately predict the qualitative patterns observed in genome wide simulations (see Figure 3 in the main text).

Figure A3 explores the parameter combinations for which we expect a transition to POD. We find that like recombination, selfing can efficiently prevent the occurrence of POD. The reason is that purging of recessive mutations is more efficient under partial or full selfing, thus increasing the frequency of the unloaded haplotype $P_{0}$.

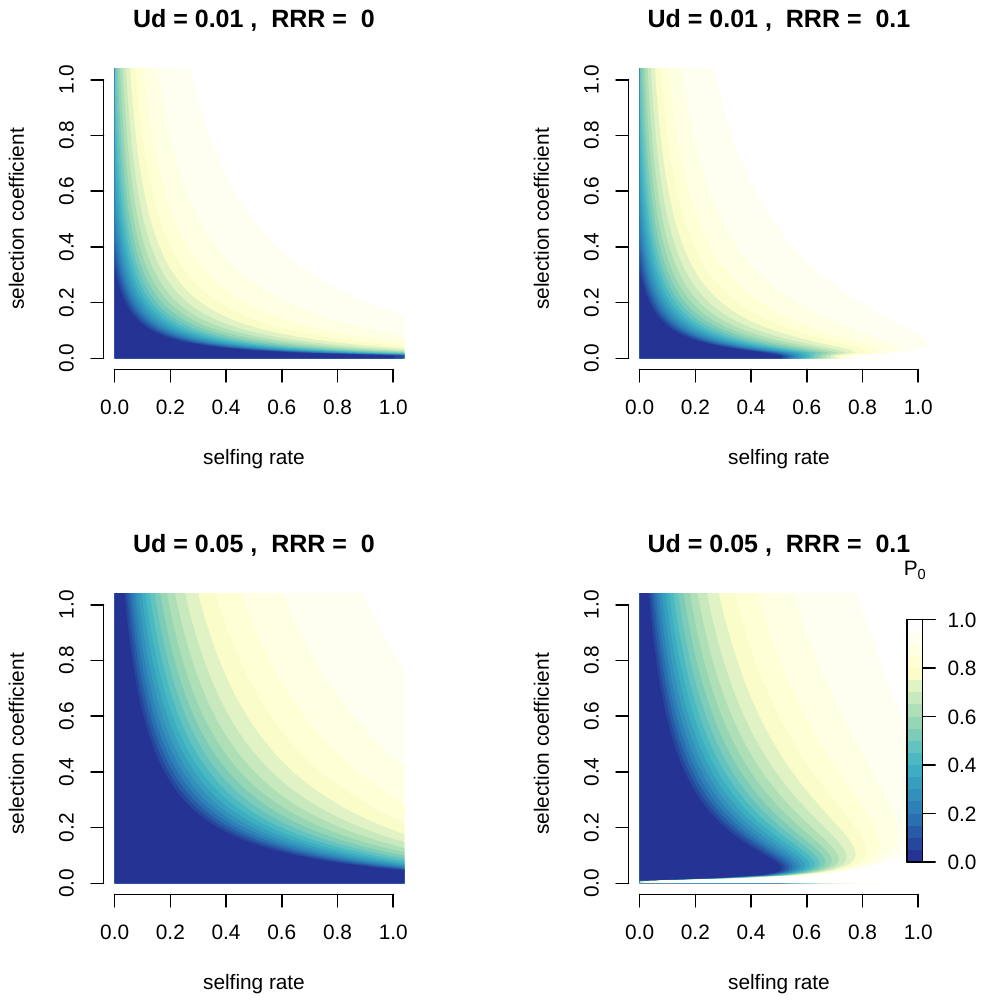

**Figure A3.** Frequency of the unloaded haplotype as a function of the selfing rate and the selection coefficient. Parameter values are n = 100 loci, and the relative recombination rate RRR = u/r and total mutation rate Ud = u*n as specified in the figure legend.

### ***Efficacy of selection with pseudo-overdominance (POD)***

We calcualte the effect of POD, that is stabilizing selection that maintains complementary haplotpes, on the efficacy of selection at linked sites. For this we assume POD occurring in a two-locus system without recombination. We consider three biallelic loci. For simplicity, we only consider the following three haplotypes: $Abc,aBc,aBC$, where capital letters indicate a derived mutation and each position correspond to a locus. Alleles $A$ and $B$ a recessive deleterious with selection coefficient $s_{1}\in[-1,0]$. The allele $C$ has selection coefficient $s_{2}>-1$ and is co-dominant.

We denote the frequencies of the haplotype $Abc,aBc$ and $aBC$ by $p_{1},p_{2}$ and $p_{3}$, respectively. We assume that the system is at POD equilibrium with $\hat{p}_{1}=\hat{p}_{2}=1/2$ and allele $C$ is introduced at low frequency on the $aB$ background.

Genotype frequencies after one generation of selfing are given by

$$p_{ii}=(1-F)p_{i}+Fp_{i}$$

and

$$p_{ij}=(1-F)2p_{i}p_{j}$$

for $i\neq j$, where $i, j=\left\{ 1,2,3 \right\}$ denote the haplotypes $Abc,aBc$ and $aBC,$ respectively. The change in genotype frequencies after selection is then given by

$$\frac{dp_{ij}}{d_{t}}=(W_{ij}-\overline{W})p_{ij},$$

where $W_{i}j$ is the fitness of genotype $ij$ and $\overline{W}$ is the mean fitness. We can simplify this system of ordinary differential equations by following the change in allele frequencies instead:

$$\frac{dp_{i}}{d_{t}}=\frac{dp_{ii}}{d_{t}}+\sum_{j\neq i} \frac{1}{2}\frac{dp_{ij}}{d_{t}}.$$

We further reduce the dimension of the system by setting $p_{1}=1-p_{2}-p_{3}$. To study the fate of the mutation $C$ linked to the $aB$ haplotype we calculate the rate of increase of the $aBC$ haplotype introduced close to the POD equilibrium $\hat{p}_{1}=\hat{p}_{2}=1/2$. We calculate the Jacobian of the system, that is the matrix with the entries $\frac{dp_{ij}}{d_{t}}$ at position $i,j$. The eigenvalues of this matrix evaluated at $p_{1}=p_{2}=1/2$ give us the rate of change of the frequency of haplotype $aBC$ when it is introduced at infinitesimal frequency. The relevant eigenvalue is given by

$$\lambda(s_{1},s_{2})=-\frac{1}{2}s_{2}(F(2(s_{1}-1)s_{2}+3s_{1}-2)+s_{1}-2).$$

Our goal is to compare the rate of change of the mutation $C$ in the case of POD to the case without POD. We can do this by calculating the ratio $eff_{sel}=\lambda(s_{1},s_{2})/\lambda(0,s_{2}).$ If this number is 1, then the initial rate of frequency change of the mutation is not affected POD. If it is smaller than 1, the efficacy of selection is reduced by the presence of POD at linked loci. We find that

$$eff_{sel}=1-s_{1}[1-\frac{1-F}{2(Fs_{2}+1)+1)}]. (A5)$$

For $F=0$ this is equal to $1-s_{1}/2$ and for $F=1$ this is equal to $1-s_{1}$, thus showing that the presence of POD reduces the efficacy of selection for linked additive mutations. This effect is amplified by selfing because the deleterious effects of the recessive mutations linked to the additive mutation become amplified by selfing. Note however, that this effect is counteracted by the decreases likelihood of observing POD in selfing populations (see Figure A3).
